# Engineering the glioblastoma microenvironment using TLR7/8 agonist-complexed graphene oxide nanosheets

**DOI:** 10.1101/2023.09.18.558196

**Authors:** Maria Stylianou, Thomas Kisby, Despoina Despotopoulou, Helen Parker, Alexandra Thawley, Kiana Arashvand, Neus Lozano, Andrew S. MacDonald, Kostas Kostarelos

## Abstract

The glioblastoma (GBM) microenvironment is characterised as immunologically ‘cold’, with immunosuppressive components that compromise the efficacy of current immunotherapies. Tumour associated macrophages and microglia (TAMMs) that are activated towards an immunosuppressive, pro-tumoral state have been identified as major contributing factors to the ‘coldness’ of GBM, while further promoting tumour progression and resistance to therapy. Based on this understanding, strategies such as macrophage reprogramming have been explored but have so far been limited by poor delivery and retention of reprogramming agents to the target cell populations within the GBM microenvironment. Consequently, clinical efficacy of such approaches has thus far shown limited success. Two-dimensional, graphene oxide (GO) nanosheets have been demonstrated to spread readily throughout the entire tumour microenvironment following a single intratumoral injection, interacting primarily with TAMMs. The current study aimed to investigate whether the immunosuppressive character of TAMMs in GBM can be ameliorated using GO sheets as a vector system to selectively deliver a TLR7/8 agonist (Resiquimod, R848), into these populations. GO enhanced the activity of R848 and induced the expression of M1-like markers on bone marrow derived macrophages *in vitro*. Using multi-parameter flow cytometry and histological analysis in a syngeneic, orthotopic mouse model of GBM, we observed that a single intratumoral injection of GO:R848 complex significantly elevated the proportion of macrophages and microglia expressing MHCII, TNFα and CD86 (associated with a pro-inflammatory, anti-tumoral state), while downregulating their expression of the M2 markers ARG1 and YM1 (associated with an anti-inflammatory, pro-tumoral state). This local complex administration inhibited tumour progression and significantly reduced tumour burden. These data illustrate that immunomodulatory GO nanosheets can effectively alter the immune landscape of GBM and modulate the wider GBM microenvironment.

**ToC Image:** 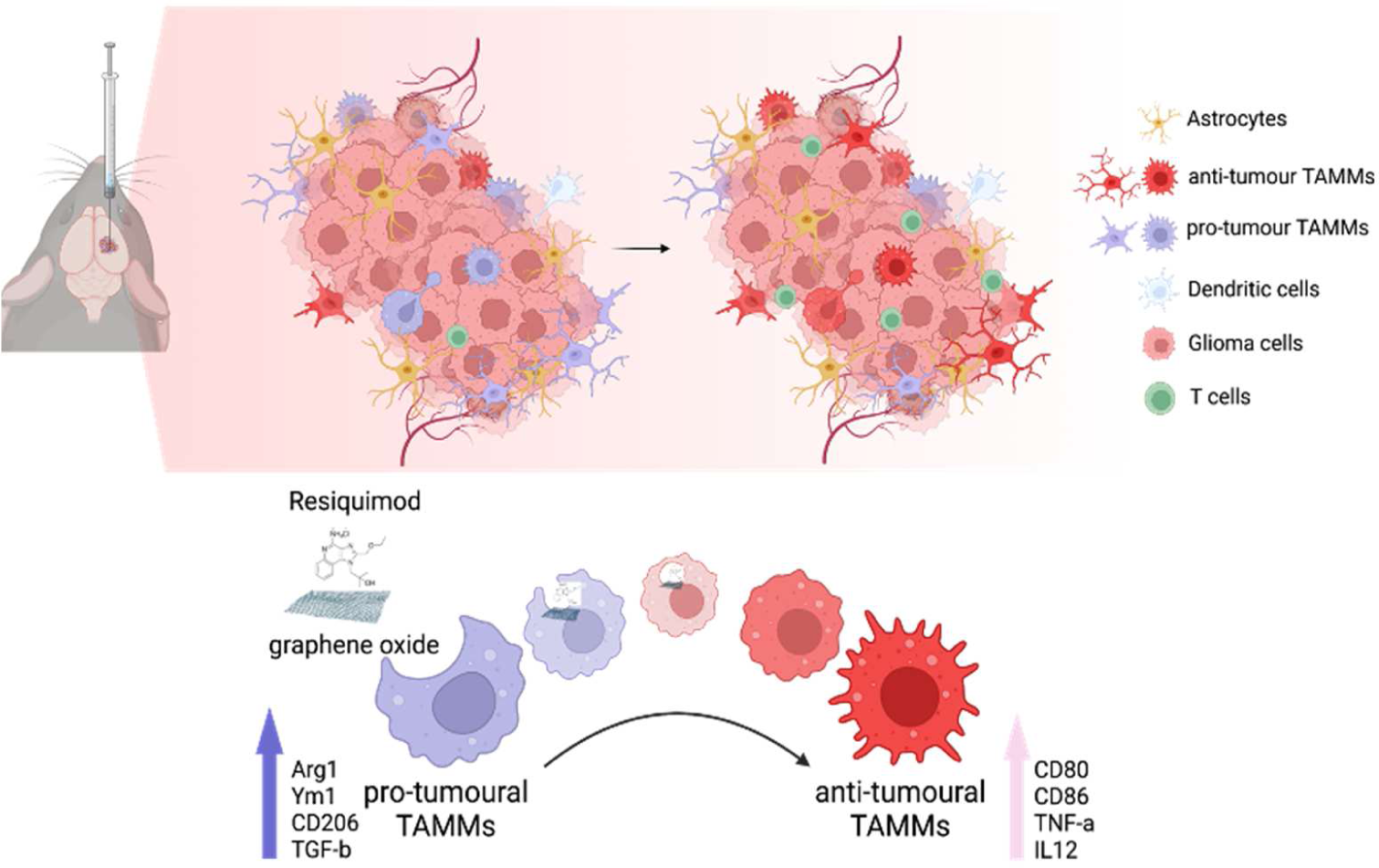

## Introduction

The tumour microenvironment (TME) plays a key role in glioblastoma (GBM) invasiveness, resistance, and progression. Despite tremendous effort to develop novel therapies including immunotherapies, GBM has proved challenging to treat. GBM has evolved multiple mechanisms to escape immune surveillance that compromise the efficacy of the standard of care and immunotherapies, with a current median survival of 15-16 months after diagnosis^1–3^.

Macrophages play a crucial homeostatic role in the clearance of cell debris and apoptotic cells by phagocytosis. Macrophage polarization is heterogeneous, with differential activation states between a pro-inflammatory/classic (M1-like) and an anti-inflammatory/alternative (M2-like) state. M1-like macrophages are responsible for pathogen and cell debris removal and have anti-tumorigenic properties, whereas M2-like macrophages promote tissue repair and have been shown to facilitate tumour proliferation^4–9^. In GBM, monocyte derived-macrophages recruited from bone marrow and brain resident macrophages known as microglia (tumour-associated macrophages/microglia, TAMMs) are abundant immune cells that can comprise 30-40% of the tumour^7, 10^. In the context of gliomas, abundance of M1-like anti-tumoral TAMMs have been correlated with low grade gliomas, while increased M2-like pro-tumoral TAMMs have been found in high grade gliomas and correlated with aggressiveness^11, 12^. Thus, targeted TAMM modulation, to increase M1-like anti-tumoral macrophages and reduce M2-like pro-tumoral activation state, has attracted great interest for treatment in high grade gliomas such as GBM.

Based on this understanding, strategies such as TAMM reprogramming have been explored alone or in combination with other immunotherapies and have demonstrated improvement of survival in GBM preclinical models^13, 14^. One of those strategies is the stimulation of Toll-like receptor (TLR) pathways using TLR agonists that leads to favourable changes in macrophage phenotype and increases their phagocytic and cancer cell clearance activity. Additionally, such stimulation could facilitate activation of adaptive immunity and support cytotoxic CD8+ T –cell responses^15, 16^. One example is the engagement of TLR7 by single stranded-RNA (ssRNA) viruses that activates NF-κB, leading to transcription of pro-inflammatory genes in various innate immune cells, including macrophages, microglia and dendritic cells^17^. TLR7 agonists include native ssRNAs and small molecules such as imidazoquinolines. The imidazoquinoline family includes the Food and Drug Administration (FDA) – approved imiquimod and its effective counterpart resiquimod (R848), with 10-fold greater pro-inflammatory activity^18^. Although there are limited studies that have introduced R848 intracranially in GBM^19–21^, it has been shown to have a potent immunomodulatory activity and exert an antitumor effect in skin malignancies, including melanoma and squamous cell carcinomas^18, 22^. Furthermore, R848 appears to modulate TAMMs from an M2-like pro-tumoral to an M1-like anti-tumoral phenotype in multiple solid tumours^23–25^. However, due to acute systemic immunotoxicities and limited capacity of many immunotherapeutics to bypass the blood brain barrier, TAMM modulation strategies have so far failed to show any clinical improvement for GBM patients^26, 27^.

The use of nanoparticles to deliver drugs directly to the tumour site can not only prolong release of the drug locally, but can also improve TAMM targeting and reduce off-target systemic side effects^28–31^. In other types of solid tumours, R848 encapsulated within nanoparticles has been demonstrated to be a potent driver of the M1-like anti-tumoral phenotype and tumour associated macrophage activation^32–34^. For instance, in breast cancer it has been demonstrated that β-cyclodextrin nanoparticles encapsulating R848 lead to significantly increased pro-inflammatory cytokines in tumour-associated macrophages, with a change toward an M1-like phenotype^24^. Graphene oxide (GO) nanosheets could be a potent platform for transport and presentation of immunomodulatory drugs in GBM, considering the large available surface area and proven biocompatibility^35–37^ without any neural-toxicity^38, 39^. Previously, we have demonstrated that GO nanosheets diffuse spontaneously throughout the tumour area of GBM without traversing its borders into the brain after a single intratumoral injection. In addition, we have illustrated that GO is taken up primarily by IBA1+ macrophages and microglia cells^40, 41^. Others have previously used cationic polymer (PEI)-modified GO to form complexes with R848 and nucleic acids^42, 43^ to demonstrate vaccine adjuvant activity in dendritic cells and macrophages *in vitro.* We have instead engineered a simpler, thinner complex between bare, medical grade, endotoxin free GO and R848, as a possible immunomodulatory platform appropriate for *in vivo* applications^44^.

In this work, we investigated thin, GO nanosheets as a platform to selectively deliver the TLR7/8 agonist, R848, into TAMMs *in vivo* as a strategy to alter the immunosuppressive character of the GBM tumour microenvironment. We prepared and characterised GO:R848 complexes and evaluated their capability to reprogram M0-like and M2-like macrophages towards M1-like macrophages, respectively. We then utilised an orthotopic, syngeneic GL261 mouse model to investigate the distribution and the *in vivo* reprogramming of TAMMs via GO:R848. We found that a single low dose intratumoral administration of GO:R848 could elevate M1-like anti-tumoral markers, such as TNFα, and alter the GBM landscape. Additionally, we showed that repeated administration could delay tumour growth *in vivo*, providing evidence that GO:R848 can therapeutically modulate the highly immunosuppressive character of GBM.

## Results

### Characterization of a non-covalent GO:R848 nanocomplex

Complexes between GO nanosheets and the R848 small molecule (GO:R848) were prepared by orderly mixing of the two components in an aqueous suspension as described (**Table S4**)^44^ and characterised via atomic force microscopy (AFM) and scanning electron microscopy (SEM) (**Figure 1**, **Figure S1**). We studied the colloidal stability and pH variation overtime with and without 5% dextrose (injection vehicle) for 24 hr and 8 days, respectively. GO:R848 and the equivalent GO alone control remained stable overtime without any significant alteration in mean particle dimensions, surface charge or pH (7.5 pH) (**Figure S2**) confirming a stable complex system suitable for further biological investigation.

**Figure 1:**
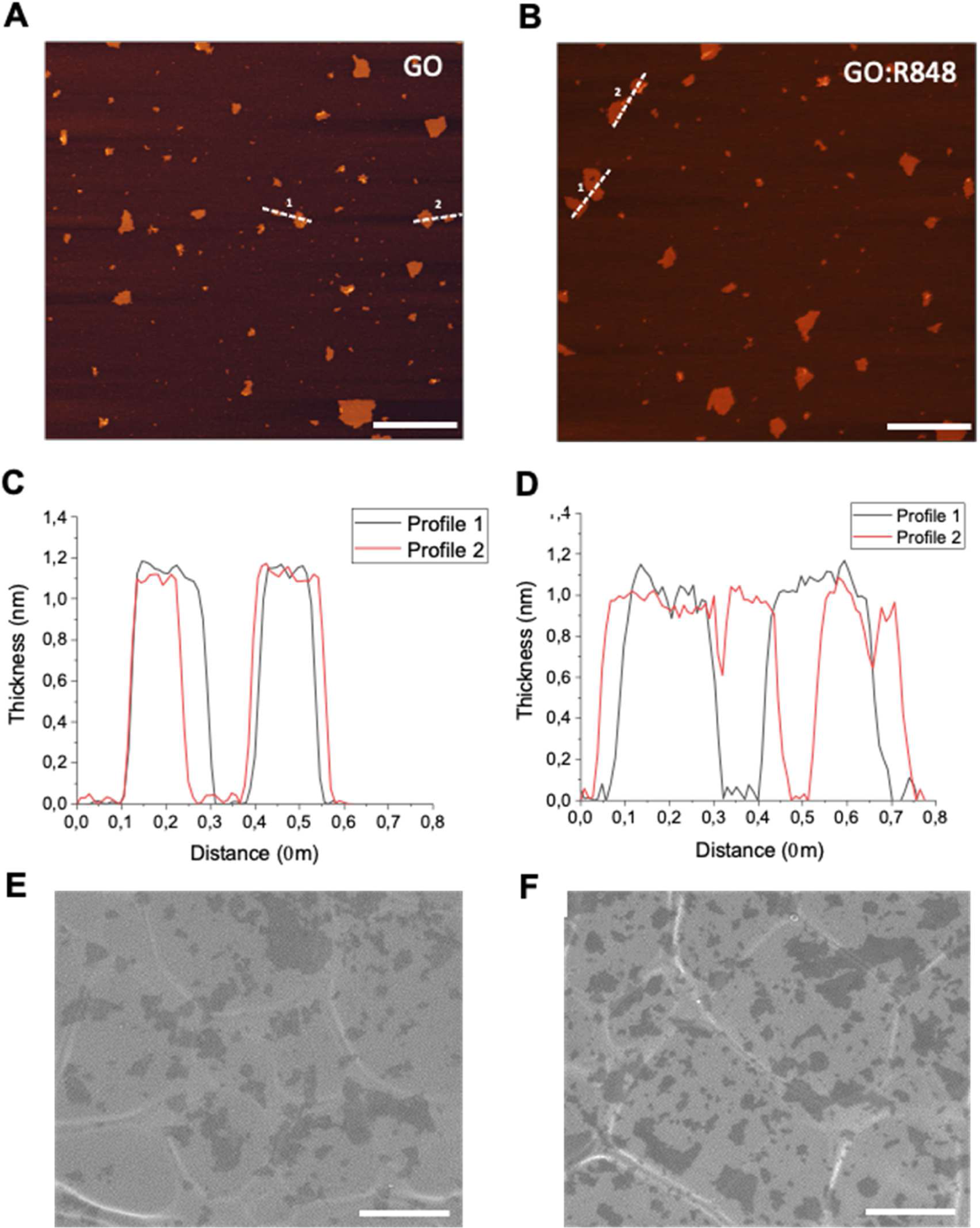
Morphological characterization of GO and GO:R848 complex. **A, B.** AFM images with corresponding flake cross-section for GO control and GO:R848. **C**, **D.** Thickness profiles based on AFM images of GO and GO:R848 n=2/graph. **E**, **F.** SEM images of GO (left panel) and GO:R848 (right panel). Scale bar, 1 μm.

### GO:R848 enhances differentiation of M0 to M1-like macrophages *in vitro*

To evaluate if GO:R848 could promote differentiation of M0 macrophages to M1-like macrophages, we first confirmed the capacity of R848 to induce TNFα production by naïve BMDMs, as a marker of M1-like activation (**Figure S3**). After treatments with GO:R848 at an R848 equivalent dose of 0.01 µg, we utilised phalloidin staining to observe macrophage morphology, as it has been highlighted as a complementary biomarker for cellular function/activation state^24, 45^. We observed via phalloidin staining that GO:R848 enhanced the differentiation of M0 to M1-like macrophages phenotypically (**Figure 2A**, **Figure S4**). We further examined this molecularly, using CD206 and CD80 membrane markers as M2-like and M1-like activation markers, respectively^46^. GO:R848 significantly elevated the proportion of CD80+ macrophages compared to M2-like (IL10/IL-4 stimulated) negative control and sustained the proportion of CD206+ macrophages at low levels, confirming an M0 to M1-like differentiation (**Figure 2B**). Consistent with this, macrophages treated with GO:R848 showed significantly elevated levels of TNFα production compared to both free R848 and the negative controls (**Figure 2C**), suggesting that the nano-complex enhances differentiation of M1-like macrophages.

**Figure 2:**
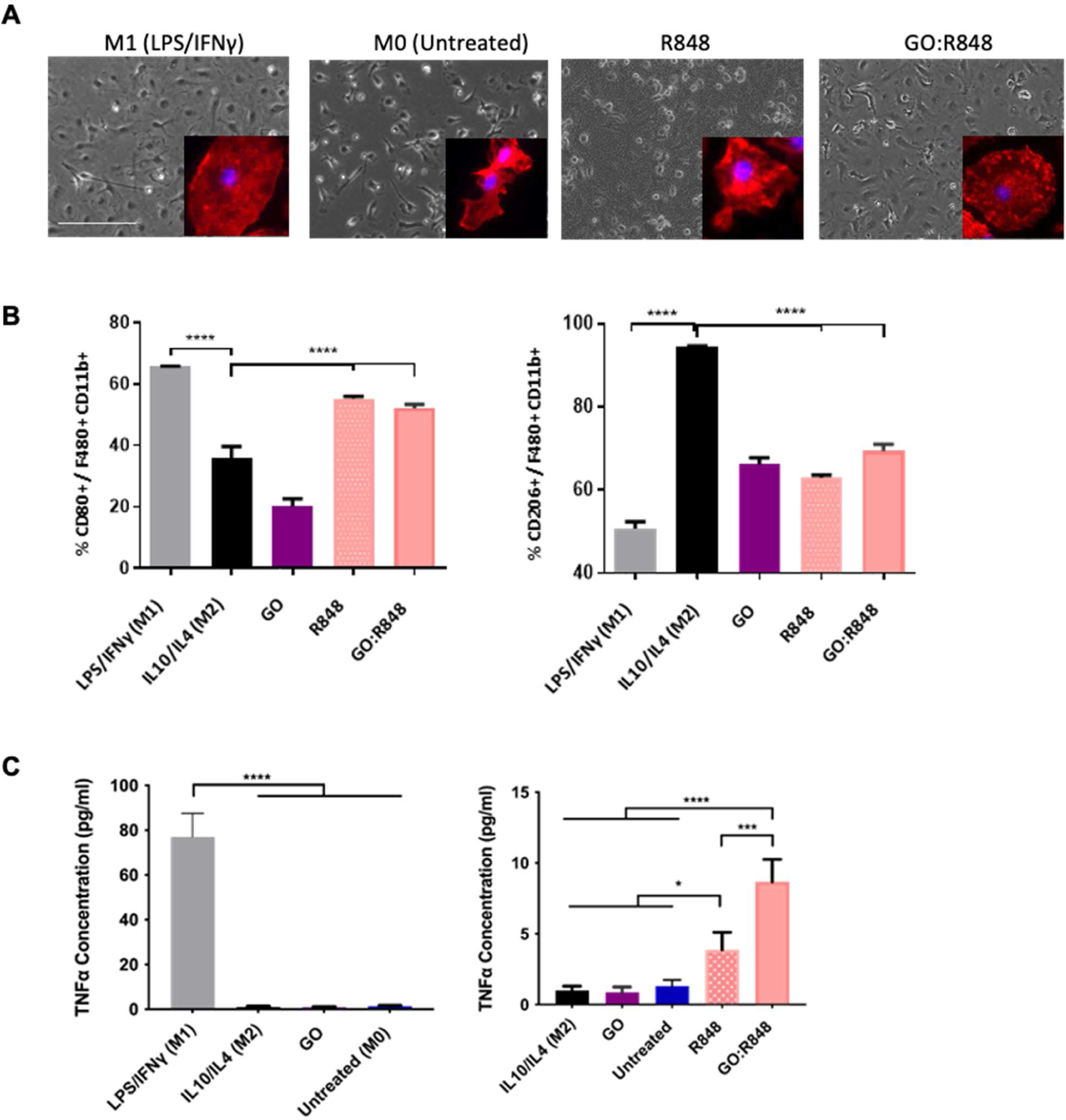
GO:R848 mediated differentiation of M0-like macrophages to M1-like macrophages. **A.** Optical images taken 24 hr post-treatment LPS (100 ng/ml)/IFNγ (20 ng/ml); M1-like anti-tumoral phenotype, IL10/L4 (20 ng/ml); M2-like pro-tumoral phenotype, GO (10 μg/ml), R848 (0.01 μg/ml) and GO:R848 (10:0.01; 10 μg: 0.01 μg). Scale bar, 400 µm. Zoomed in single cell immunofluorescence images showing single cell phenotype. Phalloidin-actin cytoskeleton; red, DAPI-nuclei; blue. **B.** Percentage of CD80+ (left side) and CD206+ (right side) out of F480+/CD11b+ cells, 24 hr post-treatment, via flow cytometry. **C.** TNFα concentration (pg/ml) from supernatant of BMDMs 24 hr post-treatment with the treatments (LPS/IFNγ, IL10/1L4, GO, R848 0.01 µg/ml and GO:R848 10:0.01). One-way ANOVA, Tukey’s multi-comparison test. (*p≤0.05, ***p≤0.001, ****p≤0.001).

### GO:R848 mediated reprogramming of M2 to M1-like macrophages *in vitro*

To examine the *in vitro* reprogramming of macrophages, we pre-treated BMDMs for 24 hr with IL10/IL4 cytokine cocktail to drive them towards an M2-like phenotype before further treatment with GO:R848 or controls. Initially, using phalloidin staining and bright field microscopy, we illustrated morphological alteration to a circular, M1-like phenotype when macrophages were treated with GO:R848 that matched the morphology of macrophages treated with LPS/IFNγ (**Figure 3A**, **Figure S5**). We observed that GO:R848, significantly attenuated the proportion of CD206+ macrophages and raised the mean fluorescence intensity of CD80+ macrophages compared to the M2-like control group, something that we did not observe with the free R848 control (**Figure 3B**, **Figure S6**). To further validate our results, we measured TNFα cytokine levels 24 hr post-treatment (48 hr post-pre differentiation to M2-like) and demonstrated that GO:R848 showed significantly advantageous M1-like cytokine production compared to free R848 (**Figure 3C**). These results suggested that GO:R848 can effectively reprogram M2-like macrophages to M1-like macrophages.

**Figure 3:**
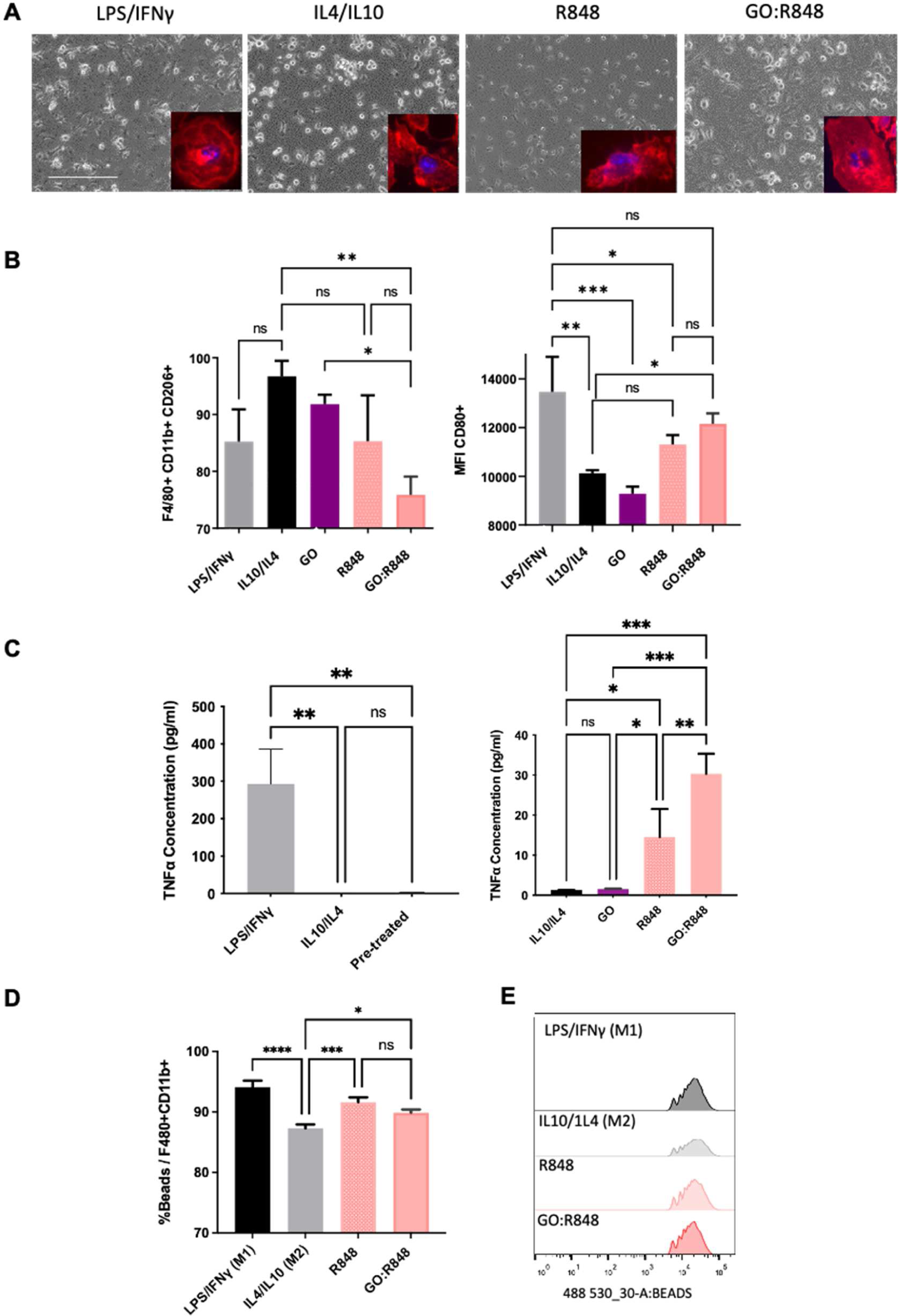
GO:R848 induced reprogramming of M2-like to M1-like macrophages with an elevated phagocytic activity. BMDMs were first pre-treated for 24 hr with IL10/IL4 to differentiate them to M2-like, followed by 24 hr of various treatments - LPS/IFNγ, IL10/1L4, GO (10 µg/ml), R848 (0.01 µg/ml) and GO:R848 (10:0.01). **A.** Optical images taken 24 hr post treatment. Scale bar, 400 µm. Zoomed in single cell immunofluorescence images showing cell phenotype. Phalloidin-actin cytoskeleton; red, DAPI-nuclei; blue**. B.** Flow cytometry charts showing the percentage of CD206+ out of F480+/CD11b+ population and the mean fluorescent intensity of CD80+ cells, 24hr post treatment. **C.** TNFα concentration (pg/ml) in supernatant of M2-like differentiated BMDMs 24 hr post-treatment (48 hours after pre-treatment) with the treatments stated above. **D.** BMDMs were pre-treated for 24 hr with IL10/IL4 (20 ng/ml) to differentiate them to M2-like macrophages, followed by 24 hr of treatment with GO:R848 or controls. Treatments were removed and cells were incubated for 1hr with green-fluorescent beads to evaluate their phagocytic ability via flow cytometry. Percentage of green-fluorescent BEADs+ cells out of the total population of F480+/CD11b+ BMDMs. **E.** Histograms illustrate the uptake and counts of fluorescent beads, 24 hr post-treatment. Statistical analysis made with one-way ANOVA, Tukey’s multi-comparison test. (*p≤0.05, **p≤0.01, ***p≤0.001, ****p≤0.001).

### GO:R848 treated macrophages retain phagocytic ability

To interrogate whether macrophages loaded with GO:R848 complex can conserve their phagocytic ability, we performed a phagocytic assay utilizing fluorescent beads. Following treatment of BMDMs with GO:R848 and controls for 24 hr, we cultured cells with fluorescent beads for 1 hr and evaluated phagocytosis via flow cytometry based on the proportion of macrophages positive for beads. Interestingly, GO:R848 treated macrophages retained their phagocytic ability in terms of proportion of bead+ macrophages and had enhanced phagocytic capacity compared to IL4/IL10 treated (M2-like) BMDMs (**Figure 3D**, **Figure 3E**). These data suggested that macrophages loaded with GO:R848 nanosheets can undergo M1-like differentiation with the elevated phagocytic activity associated with this polarized state.

### *In vivo* distribution of GO:R848 and cellular uptake

Having demonstrated *in vitro* macrophage reprogramming, we sought to investigate the *in vivo* distribution of GO:R848, after a single local administration into GBM in a syngeneic GL261 mouse model. Tumours were generated by intracranial inoculation with 5×10^4^ GL261-luc cells (day 0), followed with intratumoral (i.t.) injection of GO:R848 (10:4 mass ratio, 0.72 µg R848) on day 5 (**Figure 4A**). Histological analysis of brains on day 1, 2, 5 and 10 post-treatment (day 6, 7, 10 and 15 of tumour growth) were performed and GO:R848 complex was monitored based on the dark appearance of GO. Based on H&E staining we observed that inflammatory immune cells were recruited to the injection site and surrounded/infiltrated the area of GO:R848 complex, on day 1-post treatment (**Figure 4B**). On day 5 and 10 post-treatment, we observed the uptake of GO:R848 complex by cells and redistribution away from the injection site, spreading throughout tumour area, without traversing the tumour borders. Through further immunofluorescent analysis, we confirmed that IBA1+ TAMMs (green) were the primary carriers of GO:R848 (**Figure 4C**, **Figure S7**). These results indicated that GBM TAMMs can be passively targeted by single intra-tumoural administration of GO:R848 complexes.

**Figure 4:**
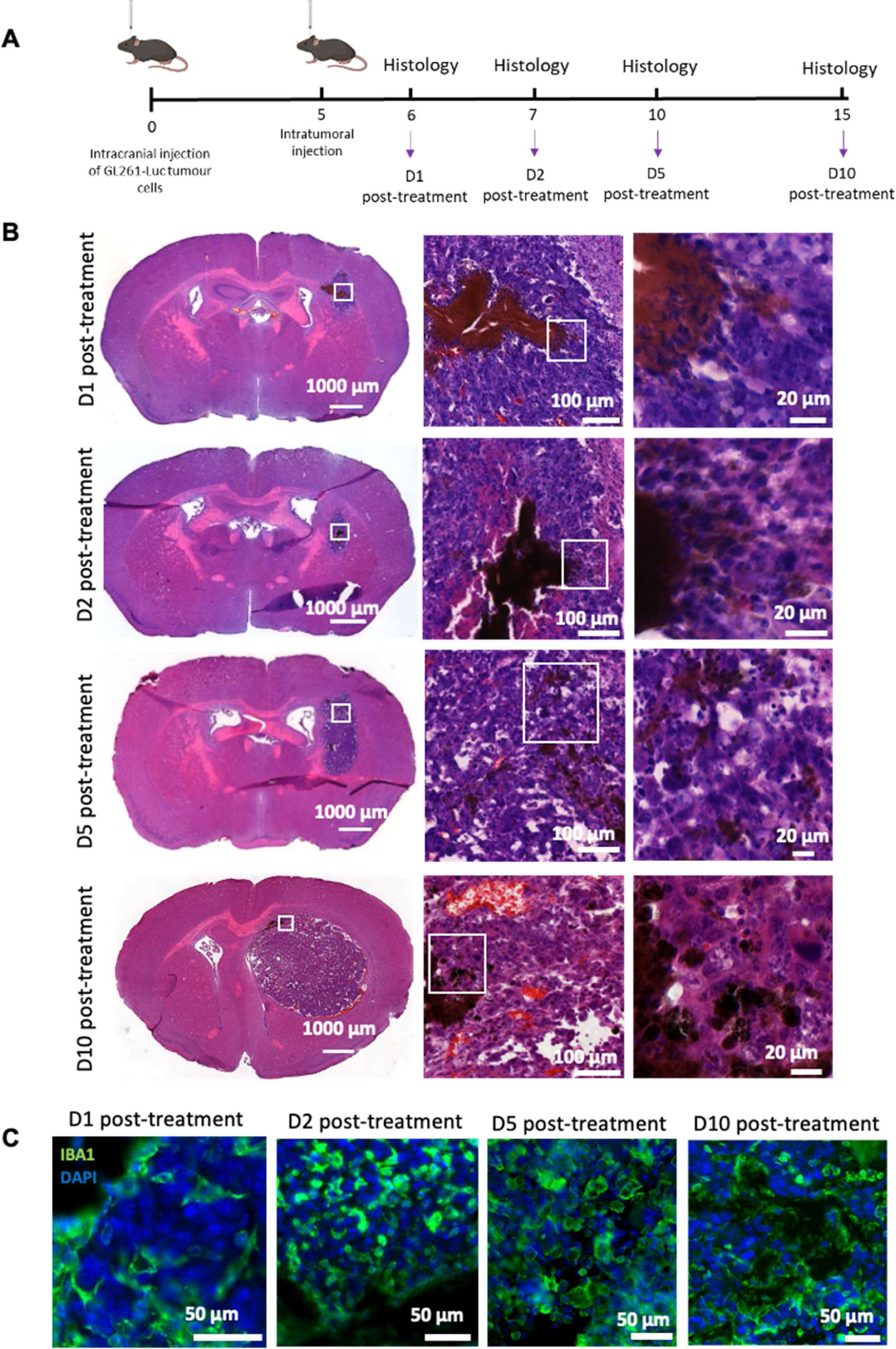
*In vivo* GO:R848 complex distribution in GBM tumours overtime. **A.** Experimental schematic illustrating the design of the *in vivo* experiment. C57Bl/6 mice were implanted with 5×10^4^ (1 µl) GL261-luc cells into the right striatum. Five days following tumour inoculation mice were treated by intratumoral (i.t.) delivery of 5% dextrose, R848 (0.72 µg) and GO:R848 (10:4; 1.8 µg: 0.72 µg) (n=4/group). **B.** Representative H&E stained sections at 1, 2 and 5 days post GO:R848 treatment. Scale bars, 1000 µm, 100 µm and 20 µm. **C.** Immunofluorescence images showing the IBA1+ cells (macrophages/microglia; green) and DAPI (nuclei; blue) around GO:R848 injected complex (black). Scale bar, 50 µm.

### *In vivo* modulation of TAMMs and alteration of GL261 microenvironment landscape

As IBA1+ TAMMs were the major cell component interacting with GO:R848, we hypothesized that we could reprogram those cells from M2-like pro-tumoral to M1-like anti-tumoral TAMMs within the TME. To evaluate the *in vivo* modulation of TAMMs in the GBM microenvironment, we generated tumours as described above, and injected GO:R848 (1.8 μg: 0.72 µg), R848 (0.72 µg) or vehicle (5% dextrose) intratumorally (**Figure 5A**). The tumour bearing brain hemispheres were collected, 24 hr and 48 hr post-treatment, and TAMMs were isolated and examined with selected activation markers via flow cytometry analysis (**Figure S8**). MHCII and CD86 activation markers were selected for the *ex vivo* evaluation of TAMMs phenotype, as they have been shown to synergistically contribute to effective antigen presentation, a key function of M1-like macrophages^47^. At both time-points, GO:R848 did not affect immune cell viability, suggesting that the GO:R848 dose used did not induce cell death *in vivo* (**Figure S9A**). As expected, 1 day post-treatment with GO:R848 or free R848, TAMMs displayed elevated expression of TNFα, MHCII and CD86 (“M1-like” anti-tumoral and activation markers) compared to vehicle control, with significantly lower mean fluorescent intensity of ARG1 (an “M2-like”, pro-tumoral marker) (**Figure 5B and 5C**). Two days post-treatment, we observed that the % of TAMMs expressing TNFα remained significantly upregulated only in the GO:R848 treated tumours compared to the controls (**Figure 5D**), suggesting that GO:R848 can reprogram TAMMs for a more sustained period of time after administration *in vivo*. Interestingly, we observed that the GO:R848 treated groups had elevated monocytes and other leukocytes 24 hr and 48 hr post-treatment, although additional analysis would be needed to determine their states (**Figure S9B and S9C**). To further validate our results, we performed immunohistological analysis to evaluate TAMM activation on days 1, 2 and 5 post-treatment (**Figure 6A**). TAMMs (IBA1+ cells) recruited to the injection site and surrounding the GO:R848 complex were a mixed population of “M2-like” (YM1+/IBA1+) and “M1-like” (CD86+/IBA1+) cells (**Figure S10**). Quantification of YM1+/IBA1+ cells showed that M2-like TAMMs were initially highly concentrated at the injection site but were significantly reduced by GO:R848 by day 2 post-treatment, compared to free R848 or vehicle treated controls (**Figure 6B, 6C and Figure S11**). In agreement, we observed that M1-like (CD86+/IBA1+) TAMMs were significantly increased throughout the tumour and injection site on day 1 and day 2 post-treatment with GO:R848, compared to the controls (**Figure 6D, 6E and Figure S12**). These data suggested that GO:R848 shifts the balance of TAMM polarization *in vivo*, decreasing M2-like TAMMs and increasing M1-like TAMMs withinin the GBM microenvironment.

**Figure 5:**
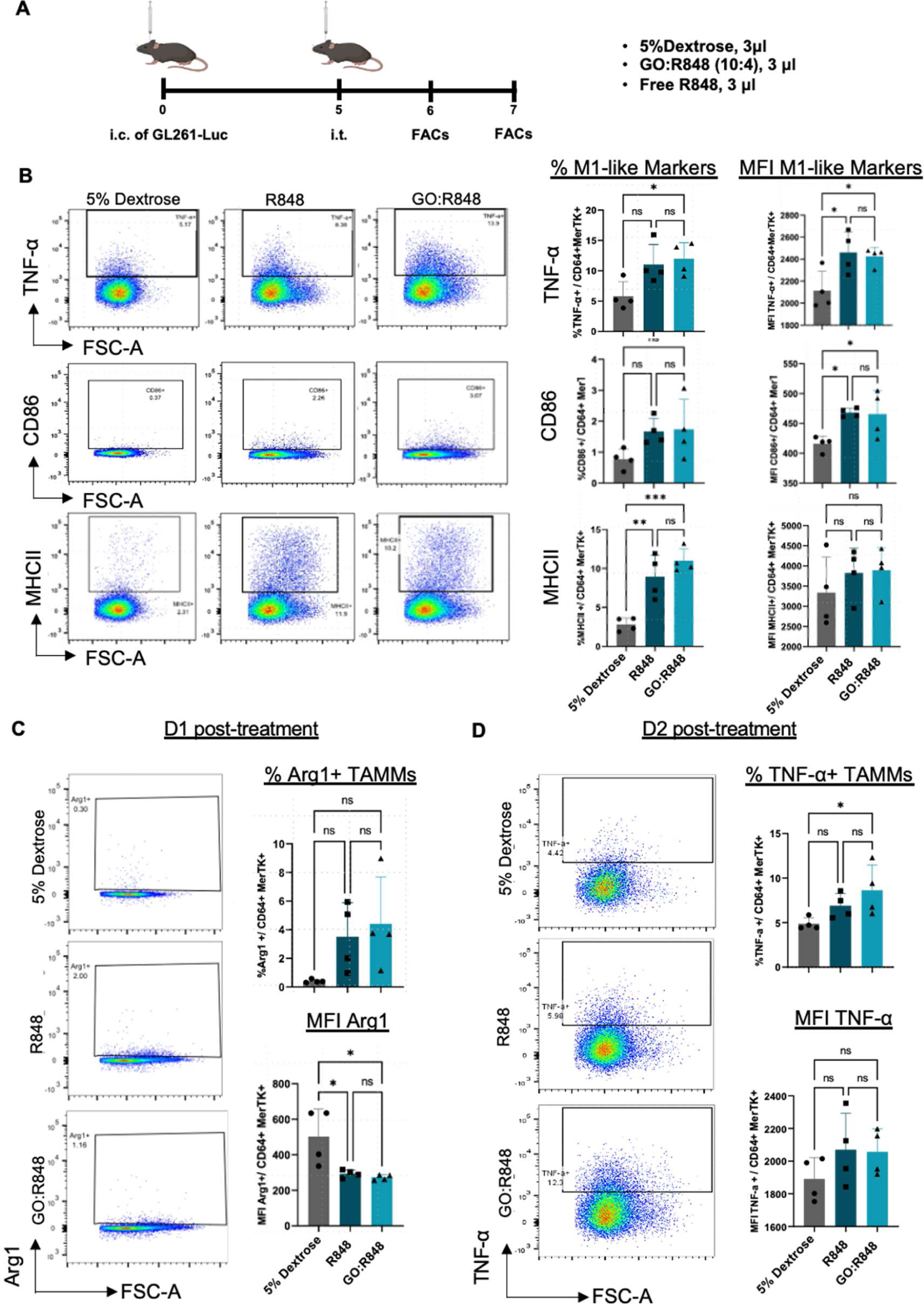
GO-R848 complex enhances M1-like anti-tumoral TAMMs, 1- and 2-days post-treatment *in vivo*. **A**. Experimental schematic illustrating the design of the in vivo experiment. C57Bl/6 mice were implanted with 5×10^4^ (1 µl) GL261-luc cells into the right striatum. BLI was conducted on day 4, as a pre-treatment baseline to randomise mice to different groups. Five days following tumour inoculation mice were treated by intratumoral (i.t.) delivery of 5% dextrose, R848 (0.72 µg) and GO:R848 (10:4; 1.8 µg: 0.72 µg) (n=4/group). **B.** Percentage and mean fluorescent intensity of pro-inflammatory cytokine (TNF-α) and activation markers (CD86 and MHCII), representative flow cytometry dot plots per treated group, on day 1 post-treatment (day 6). **C.** Percentage and mean fluorescent intensity of anti-inflammatory M2 marker (Arg1) on day 1 post-treatment (day 6), with representative flow cytometry dot plots. **D.** Percentage and mean fluorescent intensity of pro-inflammatory cytokine (TNFα), with representative flow cytometry dot plots, 2 days post-treatment. Statistical analysis made with one-way ANOVA, Tukey’s multi-comparison test. (*p≤0.05, **p≤0.01, ***p≤0.001).

**Figure 6:**
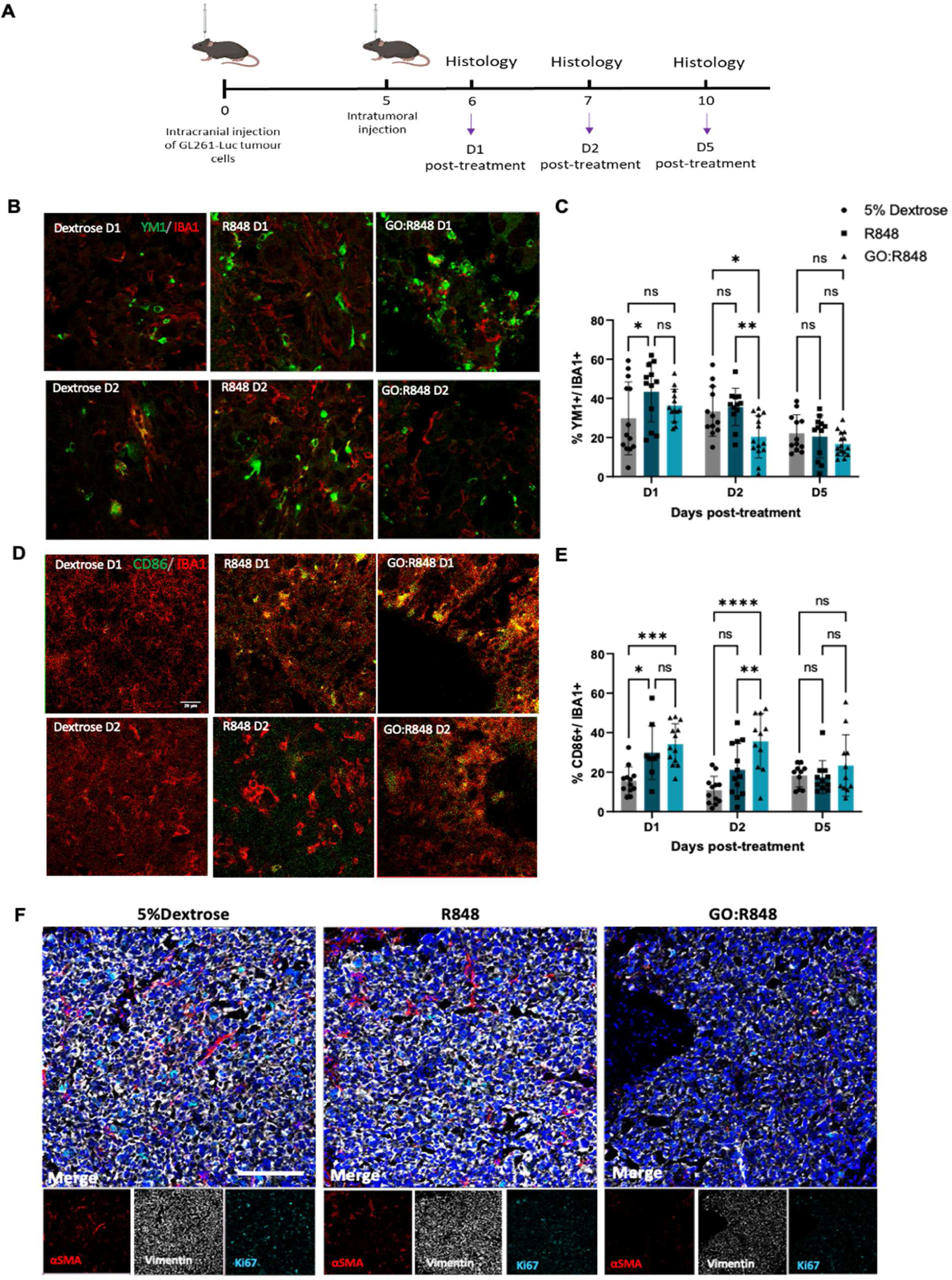
GO-R848 reduces “M2-like” pro-tumoral TAMMs and elevates the expression of M1-like anti-tumoral CD86+ TAMMs *in vivo*. **A.** Schematic illustration of the experimental design. **B.** Representative high magnification confocal images of TAMMs in GBM tissue, on day 1 and day 2 post-treatment with 5% dextrose, R848 (0.72 µg) and GO:R848 (10:4; 1.8 µg: 0.72 µg). Images show, IBA1+ TAMMs (red), YM1+ M2-like pro-tumoral TAMMs (green). Scale bar, 20µm **C.** Percentage of YM1+ cells out of the total IBA1+ cells, on day 1, 2 and 5 post-treatment with 5% dextrose, R848 and GO:R848. N=3 mice/ group. 4-5 fields of view/mouse were quantified via Image J. **D.** Confocal images of IBA1+ (red) and CD86+ (green) TAMMs 1- and 2-days post treatment. Scale bar, 20µm. **E.** Quantification of CD86+ cells out of IBA1+ cells on day 1, 2 and 5 post-treatments. Statistical analysis made with two-way ANOVA, Tukey’s multi-comparison test. (*p≤0.05, **p≤0.01, ***p≤0.001, ****p≤0.001). **F.** Representative images (600 µm x 600 µm) of tissue on day 2 post-treatment with 5% dextrose, R848 and GO:R848, showing αSMA (red);blood vessels, Vimentin (grey); epithelial to mesenchymal transition marker, Ki67 (cyan); proliferation marker, Cell mask (blue); nuclei.

Since we observed that GO:R848 reduces tumour supportive M2-like TAMMs in GBM, we hypothesized that GO:R848 could have a wider effect on the tumour microenvironment. Imaging mass cytometry (IMC) was performed on day 1, 2 and 5-post treatment to investigate several markers associated with the TME (**Figure S13A**). Remarkably, in GO:R848 treated samples, vimentin (epithelial-to-mesenchymal/GBM progression marker), Ki67 (proliferation marker) and αSMA (smooth muscle/neoangiogenesis) were reduced on day 2 post-treatment, compared to the vehicle and free R848 controls (**Figure 6F and Figures S13-S15**). Taken together, this indicates that, in addition to modulating TAMMs/immune microenvironment, GO:R848 administration induces more widespread changes to the TME.

### GO:R848 reduces tumour growth in GL261 gliomas and elevates T cell recruitment *in vivo*

Since we observed that GO:R848 treatment both modulated the polarization of TAMMs and reduced expression of tumour progression/proliferation markers, we hypothesized that GO:R848 would affect tumour growth. Intratumoral injection of GO:R848 on day 5 and day 8 post-inoculation was able to abrogate tumour growth, with mice having a significantly reduced tumour volumes, as measured by longitudinal bioluminescence imaging, up to the endpoint of the study (day 13) (**Figure 7A-C** and **Figure S16A**). The impact of GO:R848 on tumour growth was further verified via endpoint histology (**Figure 7D** and **Figure S16B**). Interestingly, this intratumoral administration of GO:R848 also increased the recruitment of both CD4+ and CD8+ T cells compared to the control, indicating further immunomodulation of the TME which may also contribute to inhibition of tumour growth (**Figure 7E, Figure S17**). Together these findings suggests that GO nanosheets non-covalently loaded with a TLR7/8 agonist can delay tumour progression through transiently reprogramming TAMMs and modulating the immunosuppressive microenvironment of GBM.

**Figure 7:**
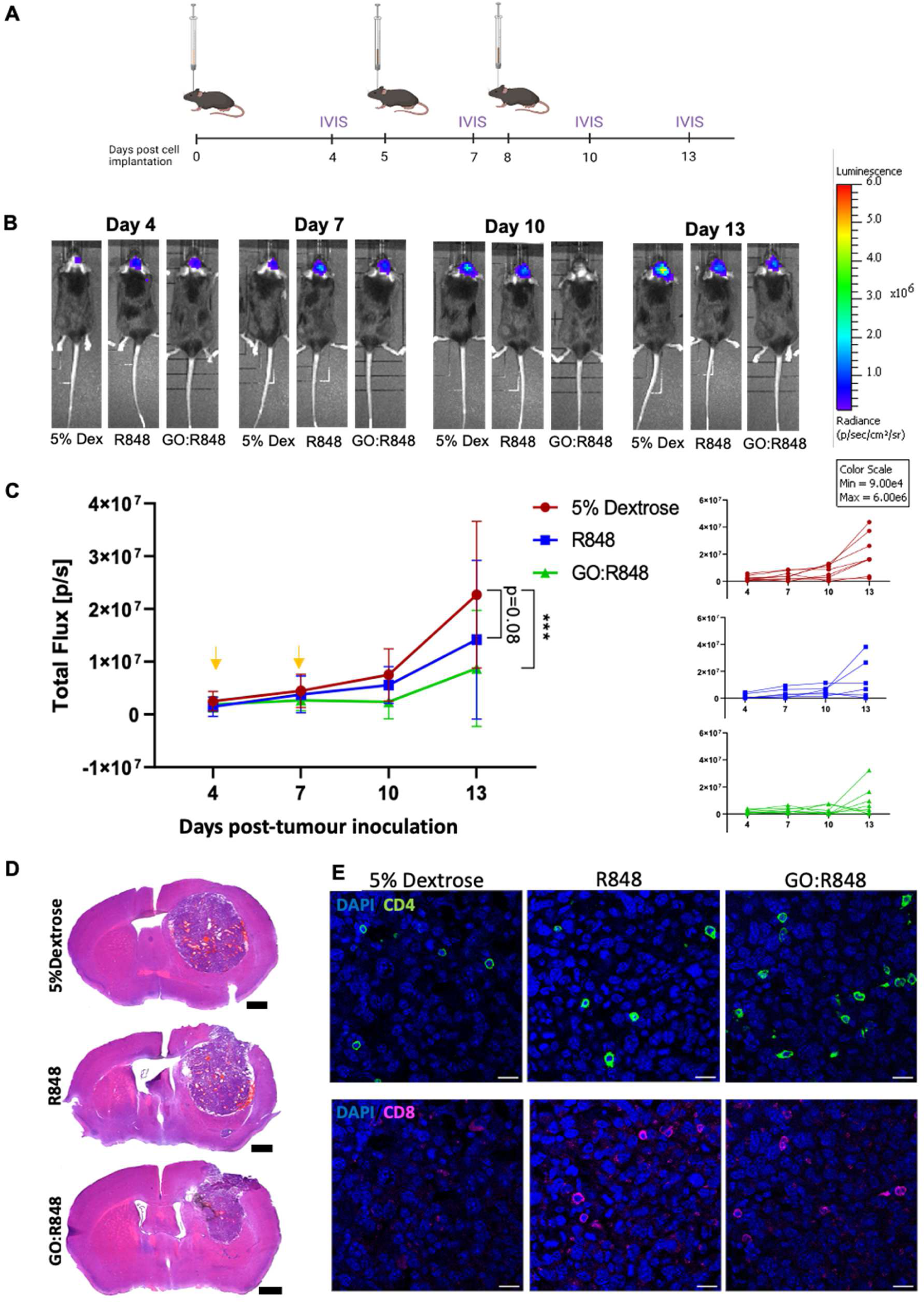
GO:R848 alters the GBM microenvironment and inhibits tumour growth. **A.** C57Bl/6 mice were implanted with 5×10^4^ (1 µl) GL261-luc cells into the right striatum. BLI was conducted on day 4, as a pre-treatment baseline to normalise mice to different groups. Five- and eight-days following tumour inoculation mice were treated by intratumoral (i.t.) delivery of 5% dextrose, free R848 and GO:R848 (10:4). Tumour growth was monitored via BLI on day 7, 9, 12 and 14 post tumour inoculation. **B.** Representative BLI images from individual mice per group. **C**. Tumour growth presented as the mean of total flux [p/s] based on bioluminescence signal per group after double intratumoral injection on day 5 and day 8 (yellow arrows) with 5% dextrose, free R848 and GO:R848 (3 µl/injection) n=7/group. Statistical analysis made with two-way ANOVA, Tukey’s multi-comparison test. (***p≤0.001). Tumour growth presented as the mean of total flux [p/s] based on bioluminescence signal per group and individual mice tumour growth for 5% dextrose, free R848 and GO:R848. **D**. Representative H&E-stained sections on day 14 post-tumour implantation, after double administration of 5% dextrose, R848, GO:R848 treatment. Scale bars, 1000 µm. **E**. High magnification confocal images of CD8+ T cells (magenta), CD4+ T cells (green) and nuclei/DAPI (blue), 14 days post tumour inoculation and post-double-treatment with 5% dextrose, R848 (0.72 µg) and GO:R848 (10:4; 1.8 µg: 0.72 µg). N= 4 mice / group. Scale bar, 20 µm.

## Discussion

Tumour-associated macrophages have been shown to be highly abundant in the stroma of a variety of solid tumours, including GBM, and have been associated with poor clinical outcomes^11, 48–50^. Strategies to reprogram these cells to a less-immunosuppressive, anti-tumor phenotype have been found to control or inhibit tumour progression ^27, 51, 52^. Despite efforts to reprogram TAMMs in aggressive brain tumours such as GBM, challenges such as efficient and preferential delivery of immunomodulatory drugs to TAMMs remain.

Here, we used small, thin GO nanosheets to transport and selectively present an immunomodulatory small molecule TLR agonist into TAMMs within the GBM tumour microenvironment. Previous studies have demonstrated that GO can be used either systemically or locally as a biocompatible delivery platform for chemotherapeutic drugs or as a vaccine adjuvant system^41, 42^. However, the application of GO for localized immunomodulation of the tumour microenvironment has not been widely investigated. A single low volume administration of GO in mouse glioma models has been evidenced to translocate throughout and persist in the TME primarily within TAMMs ^40, 41^. Here we exploited this advantageous biodistribution profile to passively target a small molecule TLR7/8 agonist (R848) into these tumour myeloid cells for the purpose of immunomodulation. R848 has been previously shown to m M2-like to M1-like reprogramming in solid tumours such as melanoma and pancreatic tumour^23, 25, 53^. Further to this, although systemic administration of R848 can induce an anti-tumor response, this has required high doses (10-60 µg) with regular repeat administration of the drug to achieve therapeutic benefit^54, 55^. In addition, systemic administration of high dose R848 has been associated with adverse effects, including brain edema, as a result of a high degree of peripheral inflammation^56^. In contrast, local administration of R848 can be used in more moderate dosages with no additional toxicities, but can be limited by rapid diffusion away from the site of injection, or metabolism of the free drug molecules. Combining R848 with a nanomaterial that is retained in the TME, such as the GO nanosheets we have employed, can increase the interaction of R848 with TAMMs and enhance the reprogramming of M2-like TAMMs toward M1-like TAMMs.

In addition to the direct effects of GO:R848 on macrophages and microglia, we also observed that this immunomodulatory nanocomplex was able to alter the wider GBM microenvironment and inhibit tumour progression. Our data further corroborate recent findings that intratumoral activation of TLR7/8 can alter the tumour microenvironment, increase peripheral lymphocyte infiltration into the tumour and inhibit tumour growth^24, 57–59^. The immunosuppressive microenvironment and paucity of tumour reactive lymphocytes has been identified as a limiting factor to the efficacy of immunotherapies. Current clinically used immunotherapies such as cancer vaccines, checkpoint inhibitors and CAR-T cells have demonstrated little or no efficacy for high grade gliomas^60^. For instance, the PD-1 checkpoint inhibitor Nivolumab failed to show any significant benefit in patient survival ^61, 62^, and cancer vaccines designed to induce an anti-tumor specific CD8+ T cell response were not successful in late stage clinical trials ^63^. Given the significant role TAMMs play in immunosuppression through secretion of cytokines, suppression of co-activation factors or expression of checkpoint molecules ^64–66^, all contributing to inducing T cell anergy/exhaustion^67, 68^, the effective reprogramming of TAMMs to a less immunosuppressive state may enhance the actions of systemic immunotherapies. Thus, combination of nanomaterial mediated local immunomodulation with existing systemic immunotherapies could solve the issue of immune-surveillance escape and low efficacy. Further studies are required to evaluate whether the GO:R848 platform presented here could provide additional synergistic effects in combination with systemic immunotherapies ^69^.

## Conclusion

In this study, we have demonstrated that GO nanosheets can provide a suitable nanomaterial platform to effectively enhance the reprogramming of M2-like macrophages/microglia to M1-like macrophages/microglia both *in vitro* and *in vivo.* Direct intratumoral administration of GO:R848 complexes offered effective modulation of the overall GBM tumour microenvironment, resulting in tumour growth inhibition in this highly aggressive brain cancer model. These findings highlight the potential of graphene-based immunomodulatory flat nanoconstructs as possible combinatorial treatment modalities to engineer an immunologically ‘hotte’ TME that could be adapted to enhance other systemic cancer therapies, such as checkpoint inhibitors.

## Experimental

### Reagents

Resiquimod (TLR7/8 agonist) was purchased from Invivogen, France and prepared following manufacturers’ instructions. Graphene oxide (GO) was synthesized and characterised at the Nanomedicine Group of the Catalan Institute of Nanoscience and Nanotechnology (ICN2) using a modified Hummers method as previously described^70^. The nanomaterials used in this study were confirmed endotoxin-free. Cell culture reagents and chemicals were purchased from Sigma-Aldrich (Merck, UK) unless otherwise stated.

### Preparation of non-covalent GO:R848 complexes

GO was neutralized at pH=10 using NaOH and R848 was added, previously reconstituted in water for injection, at the weight ratio GO:R848 of 10:4 for the *in vivo* experiments and 10:0.01 mass ratio for the *in vitro* experiments (GO concentration of 1mg/mL). The mixture was incubated for 30 min, at room temperature (RT) in an orbital shaker at 1 RCF and then incubated for a further 1 hr at RT without shaking. For the *in vitro* experiments, 50 μl of complex was mixed in 950 μl of BMDMs culture medium. For the *in vivo* experiments, complex was resuspended in 5% dextrose (Sigma, UK) at a final dose of 0.6 μg/μl GO 0.24 μg/μl R848.

### Physicochemical characterization of GO and GO:R848 complexes

#### Atomic Force Microscopy *(AFM)*

Mica surface (Ted Pella), cleaved with poly-L-lysine solution (Sigma-Aldrich) was used for the deposition of GO and GO:R848 samples, at GO concentrations of 100 µg/mL. The images were recorded in 5 µm × 5 µm dimensions, in air-tapping mode with the atomic force microscope Asylum MFP-3D (Oxford instrument) at the ICN2 Advanced Electronic Materials and Devices Group. Silicon probes (Ted Pella) were selected with 40 N/m nominal force and 300 kHz resonance frequency. The processing was performed with Gwyddion (version 2.57) and Origin (version b9.5.0.193) softwares.

#### Scanning electron microscopy *(SEM)*

GO and GO:R848 samples, at GO concentrations of 100µg/mL, were deposited on the Lacey C grid (Ted Pella). The images were recorded at the ICN2 Electron Microscopy Unit using a Magellan 400L field emission microscope (Oxford instruments) coupled with a secondary electrons detector Everhart-Thornley. The conditions used were 20 kV acceleration voltage and 100 pA beam current. Finally, the image was processed with Image J software (version 1.8.0) for the size distribution analysis.

Colloidal stability studies were performed at the ICN2 Molecular Spectroscopy and Optical Microscopy Facility with a Zeta-sizer Nano ZS (Malvern Instruments). GO and GO:R848 samples, at GO concentrations of 20 µg/mL, filled the capillary cells and were measured 3 times in room temperature by applying the water viscosity and refractive index. Data analysis was done with Zetasizer (version 7.12) and Origin (version b9.5.0.193) softwares. Data were expressed as mean ± standard deviation.

### Bone marrow derived macrophage isolation and maturation

BMDMs were isolated by flushing the bone marrow from fibula and tibias of C57/BL6 mice with DMEM medium complemented with 1% L-Glutamine, 1% Penicillin/Streptomycin and 10% FBS. Bone marrow cells were centrifuged at 300G, for 5 minutes at RT. Pellet was resuspended in medium with 10 ng/ml murine-colony stimulating factor (M-CSF) and were passed through 100 μm, strainer to remove any bone fractions. Bone Marrow cells were cultured in T25 flasks (Corning, UK), with 5% CO2, at 37 °C. For maturation, medium was changed every other day with complete medium complemented with M-CSF with a final concentration of 10 ng/ml, up to day 6 when macrophages’ maturation was tested using flow cytometry and/or used for the subsequent experiments.

### *In vitro* activation assay

Macrophage activation was determined via flow cytometry. BMDMs were seeded at a density of 150K cells per well in non-treated 24-well plate (Corning, UK) and treated for 24 h with the different treatments. Then supernatant was collected for ELISA, and cells were detached with 10 mM EDTA (Peproteck, UK) for 15 minutes at 4 °C. EDTA was neutralised with equal volume of BMDMs medium. Cells were then centrifuged at 300G for 5 min at RT and resuspended in 100 μl of PBS. Finally, cells were transferred to 96-well-V-bottom well plate for staining and flow cytometry analysis.

### Flow cytometry analysis of BMDMs

BMDM were harvested and washed with PBS by centrifugation at 300G, for 5 minutes at 8 °C. 45 μl of Zombie UV, live/dead staining (BioLegend, USA) was applied after 1:2000 dilution with PBS and the plate was incubated in the dark at RT for 15 minutes. The plate was re-centrifuged at the same speed, supernatant discarded, and cells were incubated with the conjugated primary antibody (F4/80, CD80, CD206, and CD11b) and Fc receptor blocker for 1 hr at 4 °C in the dark. Following incubation, the plate was centrifuged at 300G for 5 minutes at 8 °C. Two washing steps were conducted, and cells were fixated with 1% PFA for 10 minutes at RT was performed, following by two washing steps with flow buffer. Finally, the cells were resuspended in 200 μl flow buffer and stored in the dark at 4 °C until flow cytometry. Flow cytometry was performed using MCCIR FCF BD LSR Fortessa (BD Bioscience, UK).

### Phagocytosis assay with beads

BMDMs were plated in non-treated 24-well plates (Corning, UK) at a density of 200K cells/well and were treated for 24hr with GO:R848 and controls as mentioned above. FACs-based phagocytosis assays were performed to evaluate the phagocytic ability of macrophages towards green-fluorescent beads. Following BMDMs treatment, cells were washed once with PBS (Sigma, UK) and incubated with 1 μm green-fluorescent beads (Invitrogen, UK) for 1 hr with 5% CO_2_, at 37 °C. Macrophages were harvested post treatments and divided to 96-well-rounded-low-attachment-plate (Corning, UK) and processed as mentioned above for flow cytometry.

### Optical microscopy

Cells were imaged using a PrimoVert microscope (ZEISS) with a Primo Plan-ACHTOMAT 10X/0.25 Ph1 lens. Images were captured via an AxioCam ERc5s camera with ZEN light software. All conditions were kept consistent throughout the imaging process.

### Animals

All animal experiments were performed at the University of Manchester (UK), in accordance with the Animals (Scientific Procedures) Act 1986 (UK), approved by the University of Manchester Ethical Review Committee and under a UK Home Office Project License P089E2E0A. Animals were housed in groups of 4-5 within ventilated cages with *ad libitum* access to food and water. Female C57/BL6 (Envigo, UK) mice, 9-12 weeks old were allowed to acclimatise to the facility for at least one week prior to any procedure.

### Intracranial inoculation of glioma cells

Female C57/BL6 mice (8–9-week-old) were anaesthetised using isoflurane (2.5% induction and 1.8-2% maintenance in medical oxygen, at a rate of 1.5 - 2 L min^-1^) and placed on a stereotactic frame. Prior surgery, animals received 0.1 mg/kg of buprenorphine (Buprenex, Reckitt Benckiser, UK). A midline incision was performed to expose the cranium, dura was dried, and a 0.7 mm bore hole was drilled (Fine Science Tools, Canada) above the right striatum at 0.0 mm anterior and 2.3 mm lateral from bregma. A 10 µl Hamilton syringe (SYR10, Hamilton, USA) with a 26-gauge blunt needle (Hamilton, USA) was lowered to 3 mm below the cortical surface and slowly withdrawn 0.6 mm to create a pocket, with a final injection point at 2.4 mm depth. 5×10^4^ cells in 1 μl of PBS were injected slowly over 5 minutes at a rate of 0.2 µl /min. Post-injection the needle was kept in place for 3 minutes to minimize reflux and slowly withdrawn to minimize any innate injury. The skin incision was closed with 6-0 coated vicryl sutures (Ethicon, UK) and animals were allowed to recover in a heated environment.

### Intratumoral injection

Mice underwent intratumoral injection with 3 µl of 5% Dextose, free R848 or GO:R848 in 5% dextrose on days 5 post-tumour cell implantation. Mice were anesthetised and prepared for stereotaxic surgery as described above. The original incision was reopened, and a 33-gauge needle connected to a 10 µl Hamilton Neuros syringe was passed through the original bore hole to a depth of 2.2 mm to ensure the targeting of the tumour centre. 3 µl of GO:R848 complex or controls were injected over 15 minutes (0.2 µl / min). Post-injection the needle was kept in place for 3 minutes to minimize reflux and slowly withdrawn over 1-3 minutes. The skin incision was closed with 6-0 coated vicryl sutures and animals were allowed to recover in a heated environment, provided with mash food. For the 2^nd^ intratumoral injection, the same process was followed, with the needle lowered to a depth of 2 mm, assuming that tumour (Day 8) grew evenly.

### *In Vivo* Bioluminescence Imaging (BLI)

Tumour bearing mice were anaesthetised with 2% isoflurane (1.5% maintenance in medical oxygen) followed by intraperitoneal injection of 150 mg/kg mouse D-luciferin (15 mg/ml; Promega, UK) in PBS. 8 minutes post-injection, bioluminescence signals were detected using sequential imaging (15 measurements at 2-minute intervals) with an *in vivo* imaging system (IVIS Lumina II, PerkinElmer, UK). Images were analysed with Living Image software (version 4.7) (PerkinElmer, UK) and results were plotted in GraphPad Prism software (v6.01).

### Post-mortem tissue processing

At the end each experiment, tumour bearing mice were anaesthetized with 2.5% isoflurane and culled by cardiac perfusion with 20 ml of 2 mM EDTA in PBS. Brains were removed and fixed overnight at 4 °C in 4% PFA in PBS buffer for 24 hr, and later placed in 30% sucrose in PBS for at least 24 hr to ensure cryopreservation. The brains were snap frozen in cold isopentane (-40 °C to -50 °C) and coronal sections (20 µm thickness) were taken using a cryostat (Leica CM1950, Leica Biosystems, Germany).

### Analysis of tumour infiltrating immune cells by flow cytometry

Following perfusion, tumour bearing hemispheres were minced in a non-culture treated 12-well-plate (Corning, UK) and, 1 ml of Accutase (Sigma, UK) was added to each sample before being incubated at 37°C for 25 minutes. The digested tissue was then passed through a 100 µm strainer with the help of flow buffer (PBS + 2 mM EDTA, 2% FBS – flow buffer) and was centrifuged at 300 x g for 7 minutes at 8°C. Pellets were then re-suspended in 6ml 35% Percoll (Sigma-Aldrich, UK), samples were spiked with 2 ml of 70% Percoll (Sigma-Aldrich, UK) at the bottom of 35% Percoll layer by a 19-gauge needle and 1 ml of PBS was added on the top creating three clearly visible layers. Following centrifugation at 650 x g for 15 minutes at 20°C (acceleration 4/deceleration 1), the top-fat layer was removed, and the transparent layer was isolated. Then cells were washed with flow buffer and centrifuged at 300 x g for 7 minutes at 8°C. The medium was aspirated, and cells re-suspended in 1 ml flow buffer ready for cell counting. Samples in suspension were re-centrifuged 300 x g, 8°C for 5min. The cell pellet was then re-suspended to 1×106 cells/200 µl flow buffer which was then transferred to a 96-well V-bottom plate (ThermoFisher, UK). The plate was centrifuged at 300 x g for 5 minutes, 8°C and supernatant was removed. 10 µl of Zombie UV (1:2000), live/dead staining (BioLegend, USA) was applied, and the plate was incubated in the dark at RT for 15 minutes. Conjugated antibodies were added (**Table S2**) and the plate was then incubated for 30 minutes at 4°C in the dark. Two washing steps were conducted by removal of the supernatant, addition of 50 μl of flow buffer and centrifugation at 500 x g for 3 minutes at 8°C. Fixation with 1% PFA for 10 minutes at RT was performed, following by two washing steps with flow buffer. Finally, the supernatant was removed, and cells were re-suspended in 100 μl flow buffer and stored in the dark at 4°C until flow cytometry. Fluorescence was detected using the MCCIR FCF BD LSR Fortessa (BD Bioscience, UK) and data were analysed via FlowJo (v10.6.1). Fluorescent minus one (FMO) controls were applied for antibodies (anti-CX3CR1, anti-MerTK, anti-F4/80, anti-SiglecH, anti-MHCII, anti-CD3, anti-CD19, anti-NK1.1 and anti-TCRb antibodies) for the purpose of gating.

### Haematoxylin and Eosin Staining (H&E)

Sections were stained with H&E staining to observe the histological characteristics of the tumour sections and determine the tumour volume. Cryo-sections were left for 10 min at RT to dry and staining was performed as mentioned before^41^. The whole staining process was performed automatically by Leicia autostainier (Leica Biosystems, Germany). Slides were scanned using a 3D Histech Panoramic 250 slide-scanner and analysed using CaseViewer software (2.4.0.119028).

### Immunofluorescence

Cryosections were removed from the freezer and allowed to thaw prior to being post-fixed in ice-cold acetone for 10 minutes and permeabilised in 0.3% Triton-X (Sigma, UK) solution. Sections were then incubated with blocking buffer composed of 5F% (v/v) Normal Donkey Serum (Sigma, UK), 1% Bovine Serum Albumin (BSA) (Sigma, UK) in PBS-Triton X 0.2% for 1.5 hr at room temperature. Diluted primary antibodies were added on sections and incubated in a humidified chamber at 4 °C, overnight. Sections were then washed 3 times with washing buffer (0.2% Triton-X in PBS) and then incubated with diluted secondary antibodies for 1 hr at room temperature. After, 6 more washing steps with the washing buffer, 1X PBS and distilled water, sections were mounted with Prolong Gold Antifade medium containing DAPI and let dry at RT, overnight in the dark. Fluorescence microscopy images were acquired a Leica TCS SP8 AOBS inverted confocal microscope using the 63x objective with optical Z spacing as specified by the LASX imaging software (Leica, UK). The antibodies used in these investigations are given in Table S1.

### Image Analysis

Images were analysed and quantified using ImageJ (NIH, USA). Z-stack images separated to different channels and each marker was normalised based on the negative control (only secondary antibody). IBA1+ cells were manually counted via cell counter per field of view and were co-localized with CD86 or YM1 channels. Individual CD86+ or YM1+ cells were counted using cell counter function and the % of double positive cells was calculated out of the total IBA1+/field of view.

### Image Mass Cytometry (IMC) – Staining, data acquisition and analysis

Cryosections of thickness 10 µm were let to thaw prior to being post-fixed in iced cold acetone for 10 min. Sections were incubated for 10 min in warm sodium citrate buffer for antigen retrieval and buffer was let to cool down at RT, until were permeabilised in 0.3% Triton-X (Sigma, UK) PBS solution. Sections were then incubated in 3% of BSA (Sigma, UK) in PBS for 1.5 hr at RT. Metal-conjugated antibodies were diluted in 0.5% BSA, added on sections and incubated in a humidified chamber at 4°C (**Table S2**). Sections were washed 3 more times with the washing buffer and sections were incubated with Iridium (Fluidigm, US) at a 1:500 dilution factor in PBS to stain the DNA. After 3 more washing steps, sections were let dry at RT and stored at RT until tissue ablation and IMC acquisition.

The areas of acquisition were chosen based on the distinct area of the tumour and the regions of interest (ROI) were selected to ablate for each brain section. Prior acquisition, the Hyperion mass cytometry system was autotuned using a 3-element tuning slide according to the provider protocol. As an extra verification point for a successful tuning, a detection of at least 500 mean duals of 175Lu was used. The slides with sections stained with the IMC panel containing 25 metal-conjugated antibodies (**Table S3**) were placed in Hyperion (Fluidigm, US) one by one using the same laser power. The chosen ROIs were ablated and acquired at 200Hz. Data were exported as MCD files and visualised using the Fluidigm MCD viewer software. For data analysis, MCD viewer was used to set minimum and maximum threshold for each image based on the unstained control. All thresholds were kept the same for each marker across the different ROIs. Merge images were composed using ImageJ software (NIH, USA).

### Statistical analysis

Data analysis and graphical design were performed using GraphPad Prism software (v6.01). Flow cytometry data analysis was completed using FlowJo (v10.6.1). P-values were calculated using two-way or one-way ANOVA with Tukey’s post-hoc test for multiple comparisons, unless stated otherwise in figure legends. P-values of <0.05 were considered statistically significant. Data is plotted as mean ± standard deviation (SD) unless stated otherwise.

## Acknowledgements

M.S., A.T., K.A, H.P, A.M and K.K. would like to acknowledge the United Kingdom Research and Innovation (UKRI) Engineering and Physical Sciences Research Council (EPSRC) 2D-Health Programme Grant (EP/P00119X/1) for financial support. T.K. and K.K. would also like to acknowledge the UKRI (EPSRC) for an International Centre-to-Centre grant (EP/S030719/1). The ICN2 is funded by the CERCA programme, Generalitat de Catalunya, and is supported by the Severo Ochoa Centres of Excellence programme by the Spanish Research Agency (AEI, grant no. SEV-2017-0706). The authors would like to acknowledge the ICN2 Advanced Electronic Materials and Devices Group (Prof. Jose A. Garrido) for the access to the AFM instrumentation. The authors would also like to thank Kate Hills, and Abbie Dodd for their assistance to surgeries and sample processing. The authors would also like to thank the Bioimaging core facility, Biological Services facility, Flow cytometry facility and Histology core facility at the University of Manchester for technical support and access to equipment.

## Author contributions

M.S., T.K. and K.K. initiated and designed the study. K.K. and T.K. coordinated the study. M.S. conducted *in vivo*, *in vitro* experiments and data analysis. T.K., A.T. and K.A. assisted with *in vivo* experiments, *in vivo* imaging, and tissue processing. D.D. and N.L. developed the complexation protocol and D.D. performed the complex characterization. A.M. and H.P. advised in experimental design and interpretation of data. M.S., T.K., D.D., N.L., A.M. and K.K. contributed to the writing of the manuscript.

## Notes

### Competing Interest Statement

The authors have declared no competing interest.

