## Supporting Information for "Engineering the glioblastoma microenvironment using TLR7/8 agonist-complexed graphene oxide nanosheets"

---

### SUPPORTING INFORMATION

#### Experimental

**Preparation of Compensation beads for flow cytometry.** UltraComp eBeads (Thermofisher, UK) were used to compensate the different channels and avoid overlapping of spectra. One drop of beads was diluted with 1% FBS/PBS for every 5 compensation controls. 100 µl of diluted beads were mixed with each individual antibody (diluted 1:100) separately (20 µl) and incubated in dark at RT for 10 minutes. Post incubation, 1 ml of 1% FBS/PBS was added and were centrifuged at 300G for 5 minutes. Samples were re-suspended in 200 µl of 1% FBS/PBS, ready for acquisition. For Live/Dead UltraComp compensation eBeads (Thermofisher, UK), beads were warmed to RT for 5 minutes, one drop of beads and 10 µl of Amine DYE (Zombie UV) at 1:2000 dilution were mixed and incubated in dark for 30 minutes at RT. 1 ml of PBS was added and the sample was centrifuged at 300G for 5 minutes. Supernatant was aspirated and one drop of unlabelled beads was added. The sample was then diluted in 200 µl PBS, ready for acquisition.

**pH measurements.** For the pH measurements, the FiveEasy™ FP20 pH meter equipped with the Mettler Toledo™ pH Electrode InLab Ultra-Micro-ISM was used.

**Table S1:** Primary and secondary antibodies used for immunofluorescence staining of brain sections.

| Primary Abs | Secondary Abs | Target | Dilution | Purchased |
|---|---|---|---|---|
| Rabbit anti-mouse | - | IBA1 marker (wako) – Macrophages/Microglia | 1:500 | Fujifilm - 019-19741 Wako |
| Goat anti-mouse | - | IBA1 marker - Macrophages/Microglia | 1:500 | Abcam ab5076 |
| Goat anti-mouse | - | YM1/Chitinase 3-like 3 Biotinylated Antibody | 1:100 | R&D systems-BAF2446 |
| Anti-mouse AF700 | | CD86 | 1:50 | Invitrogen 56-0862-82 |
| - | Donkey anti-goat Alexa Fluor 647 | Goat | 1:500 | Invitrogen A21447 |
| **-** | Donkey anti-rabbit Alexa Fluor 488 | Rabbit | 1:500 | Invitrogen A21206 |

**Table S2:** Conjugated antibodies used for multi-color flow cytometry

| Channels | Marker | Fluorochrome | Dilution factor | Cat No. |
|---|---|---|---|---|
| **1** | CD163 | FITC | 200 | Invitrogen 11-1631-80 |
| **2** | CX3CR1 | PerCP/eF710 | 50 | Biolegend 149010 |
| **3** | MerTK | A647 | 50 | Biolegend 151508 |
| **4** | CD86 | AF700 | 100 | Invitrogen 56-0862-82 |
| **5** | Arg1 | BV421 | 100 | Invitrogen 48-3697-82 |
| **6** | Ly6C | BV510 | 600 | Biolegend 128033 |
| **7** | CD45 | BV605 | 200 | BD Bioscience 563051 |
| **8** | CD11c | BV650 | 200 | BD Bioscience |
| **9** | CD11b | BV711 | 500 | Biolegend 101228 |
| **10** | TNFa | BV785 | 160 | Biolegend 506341 |
| **11** | Zombie UV L/D | Live/Dead Blue | 2000 | Biolegend 423108 |
| **12** | Siglec H | PE | 50 | BD Bioscience 565527 |
| **13** | CD49d | PE/CF594 | 200 | BD Bioscience 564395 |
| **14** | MHCII | PE/Cy5 | 60 | Biolegend 107612 |
| **17** | CD64 | PE/Cy7 | 100 | Biolegend 139314 |
| **18** | CD3<br>CD19<br>NK1.1<br>TCRb<br>Siglec F<br>Ly6G | APC/eFluor780 | 100<br>100<br>200<br>200<br>150<br>80 | Invitrogen 232587<br>Invitrogen 47-5941-82<br>Invitrogen 47-0193-82<br>Invitrogen 47-5961-82<br>BD Bioscience 502681<br>Invitrogen 47-9668-82 |

**Table S3:** Antibodies and Lanthanides used for IMC analysis

| Channels | Marker | Lanthanide | Dilution factor | Cat No. |
|---|---|---|---|---|
| **1** | Alpha-SMA | 141Pr | 1:200 | Abcam ab215368 |
| **3** | Vimentin | 147Sm | 1:100 | Abcam ab223871 |
| **4** | CD45 | 152Sm | 1:50 | Thermo Scientific 14-0452-82 |
| **5** | CD64 | 153Eu | 1:50 | Thermo Scientific MA5-29705 |
| **6** | E-cadherin | 158Gd | 1:100 | Abcam ab239883 |
| **7** | NF-kB | 166Er | 1:400 | Abcam ab207297 |
| **8** | CD8a | 169Tm | 1:100 | Thermo Scientific 14-0808-82 |
| **9** | CD3e | 170Er | 1:100 | Biolegend 362701 |
| **10** | Ki-67 | 172Yb | 1:100 | Abcam ab197547 |
| **11** | CD4 | 175Lu | 1:100 | Thermo Scientific 14-9766-82 |
| **12** | TMEM119 | 171Er | 1:150 | Abcam ab234501 |
| **13** | Iridium (DNA mask) | Ir191/193 | 1:400 | Fluidigm |

**Table S4:** Characterization of GO and GO:R848 complex. Physicochemical properties including degree of defects, interlayer distance, functional groups and chemical composition identified by different techniques are presented. This Table has been adapted from Despotopoulou et al.[44] where the complete characterization of the GO:R848 complexes has been presented.

| Physicochemical properties* | Technique | GO | GO:R848 |
|---|---|---|---|
| Degree of defects ($I_D/I_G$) | Raman Spectroscopy | 1.24 | 1.11 |
| Peak (2θ) | XRD | 11.59 ° | 10.02 ° |
| Interlayer distance (nm) | | 0.76 | 0.88 |
| Functional groups | FTIR | v(C=O): 1726 $cm^{-1}$<br>v(C=C): 1625 $cm^{-1}$<br>v(O-H): 1400 $cm^{-1}$<br>v(C-O): 1046 $cm^{-1}$ | v(C=O): 1726 $cm^{-1}$<br>v(C=N): 1678 $cm^{-1}$<br>v(C=C): 1616 $cm^{-1}$<br>v(O-H): 1395 $cm^{-1}$<br>v(C-O): 1041 $cm^{-1}$ |
| Chemical composition (%) | XPS | C: 61.7, O: 29.9, Na: 4.6, Cl: 0.6, S: 0.5, Si: 2.7 | C: 59.3, O: 32.1, Na: 4.0, N:1.2, Cl: 0.8, S: 0.4, Si: 2.2 |

### Supporting Figure 1

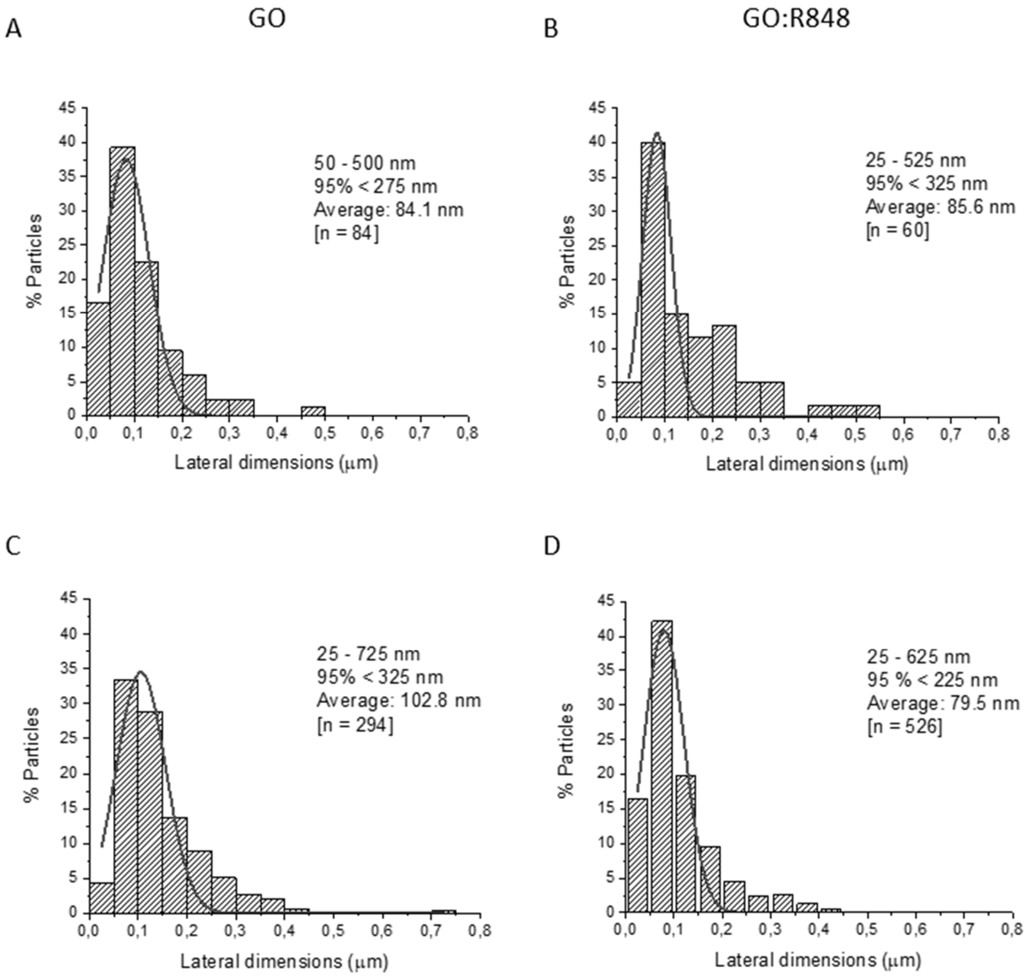


**Figure S1: Size distributions of GO and GO:R848.** Based on AFM images (**A** and **B**) and corresponding size distributions based on SEM images (**C** and **D**) for GO alone and GO:R848 complexes.

#### Supporting Figure 2

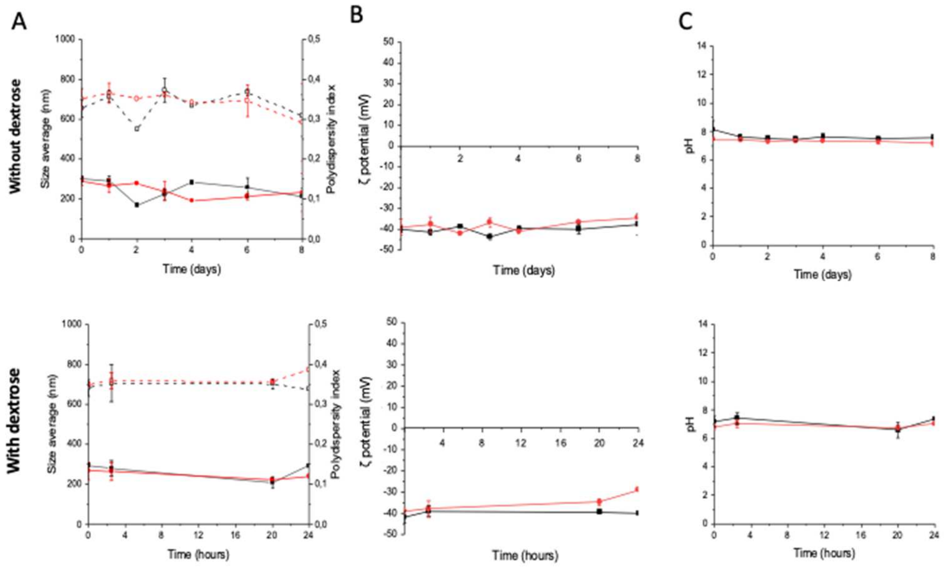


**Figure S2. Colloidal stability over time for the GO alone control (black) and the GO:R848 complex (red) with and without 5% dextrose. A.** Mean particle size (solid symbols) and polydispersity index (open symbols) measurements at different time points by dynamic light scattering (DLS). **B.** Zeta-potential measurements; and **C.** pH measurements. At least n=3 replicates were measured for each condition.

#### Supporting Figure 3

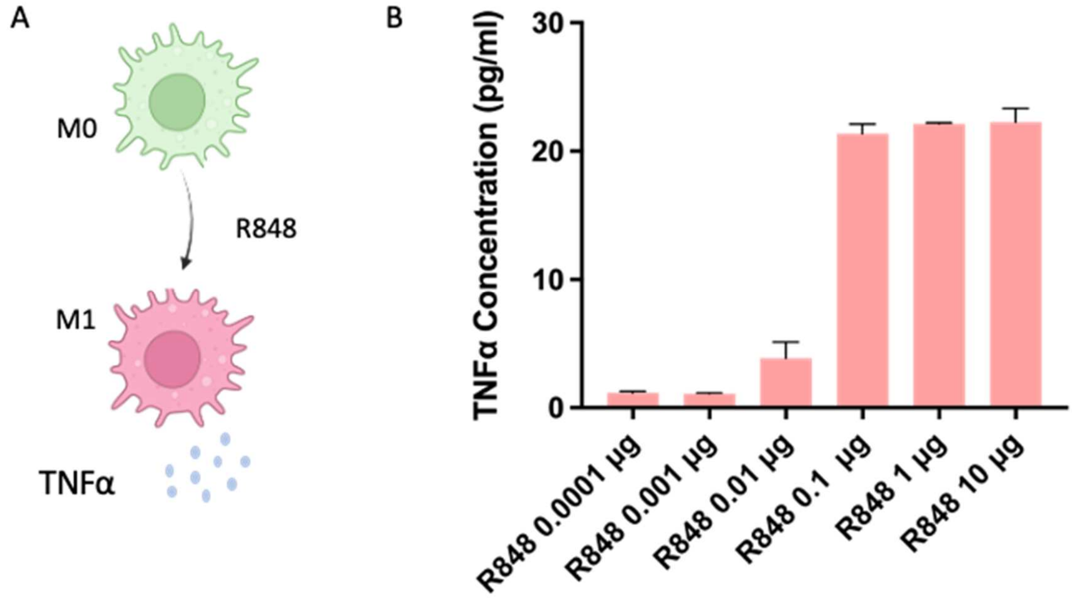


**Figure S3: Dose response of M1 differentiation mediated by R848. A.** Schematic of M0 to M1 differentiation**. B.** Concentration-depended TNFα production (pg/ml) as an M1-like driven cytokine indicator, of BMDMs treated with R848, 24 hr post-treatment.

#### Supporting Figure 4

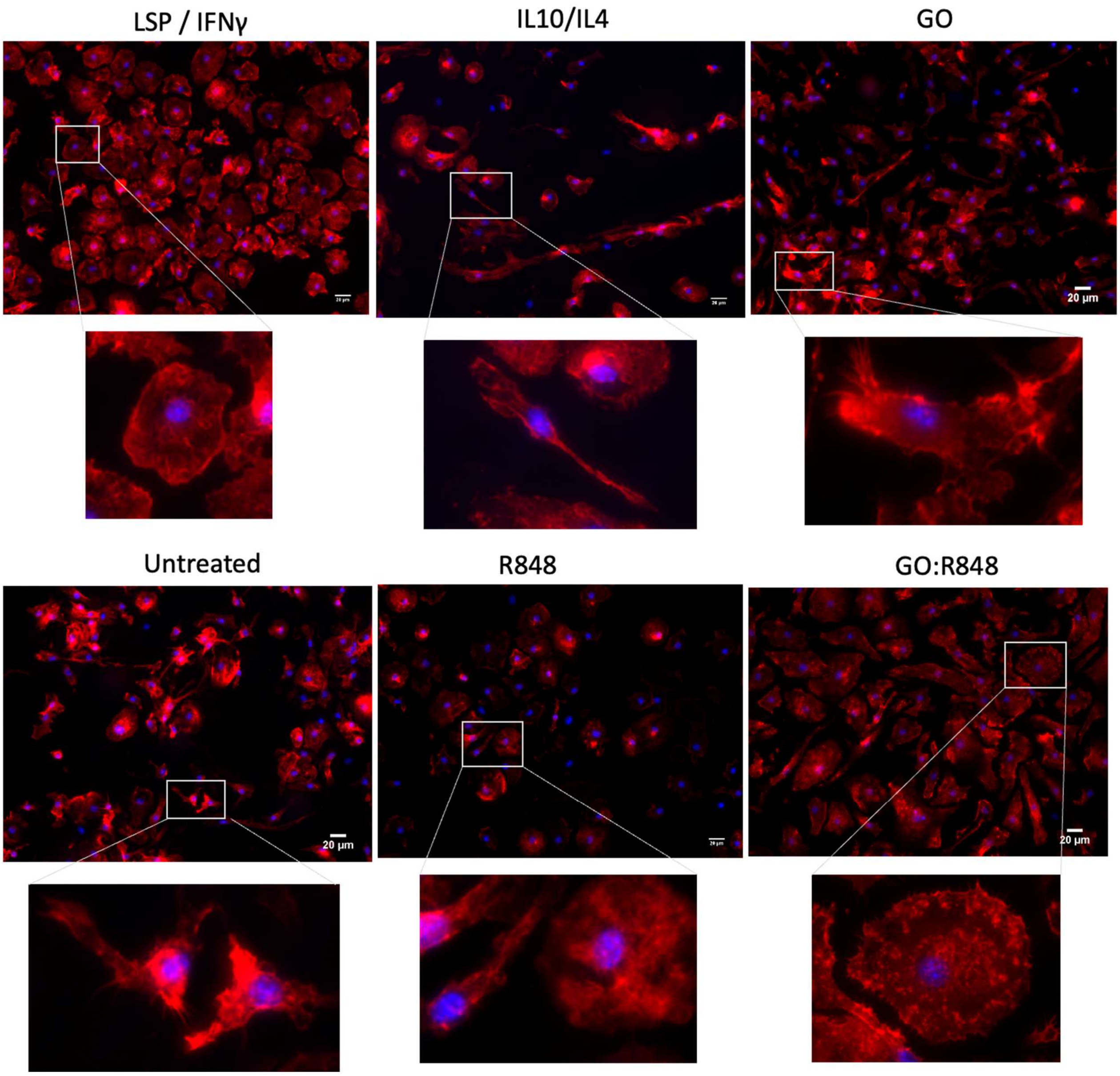


**Figure S4: Phenotypic differentiation of M0 to M1 BMDMs 24hr post- treatment with GO:R848.** BMDMs were treated with LPS/IFNγ, IL10/IL4, GO (10µg/ml), R848 (0.01µg/ml), GO:R848 (10:0.01) for 24 hr. Red staining indicates phalloidin, marker for cell actin showing cell morphology. Scale bar, 20 µm.

#### Supporting Figure 5

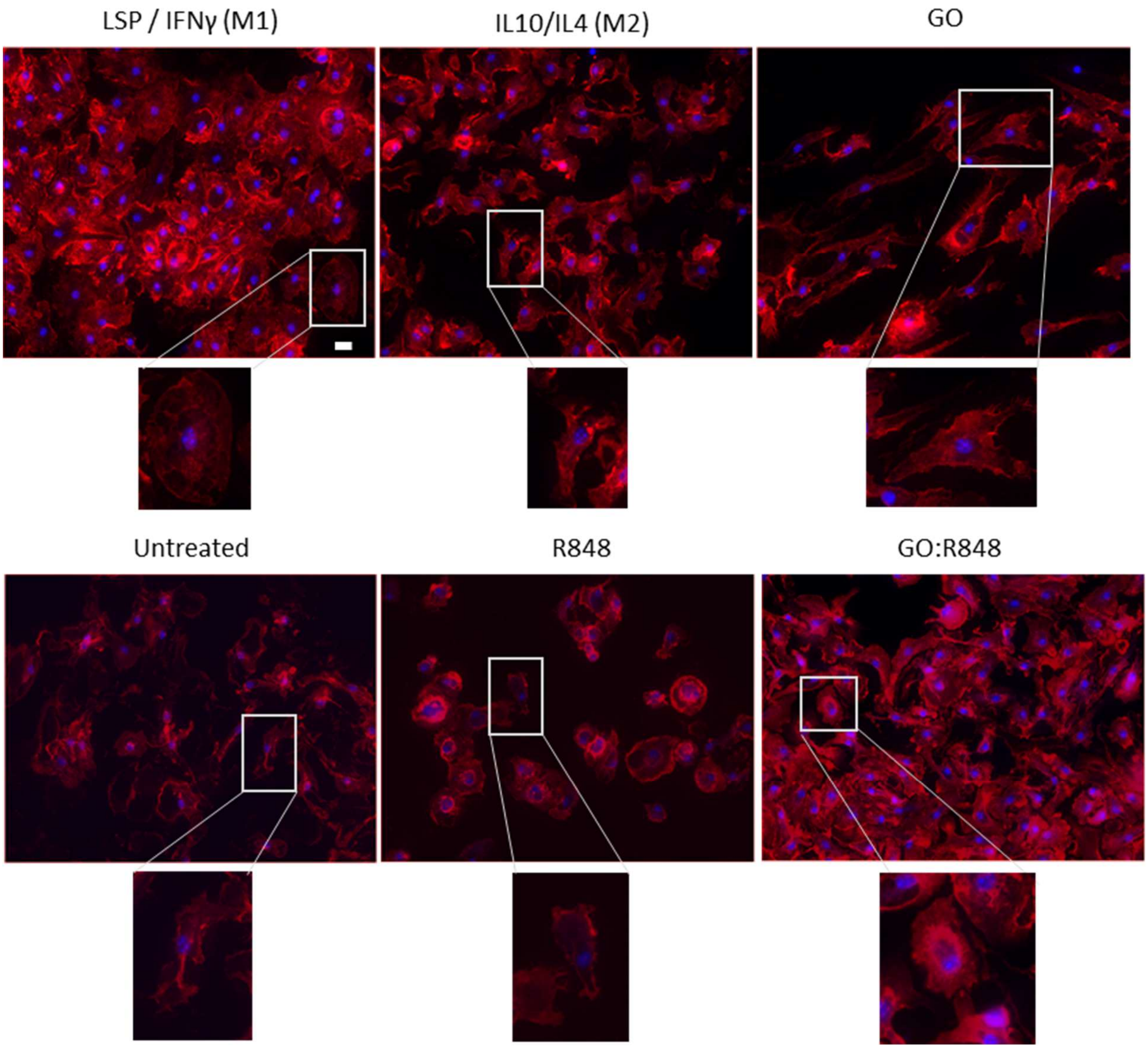


**Figure S5: Phenotypic differentiation of M2 to M1 BMDMs 24hr post- treatment with GO:R848.** BMDMs were treated with IL10/IL4 (20 ng/ml) for 24hr following with LPS/IFNg, IL10/IL4, GO (10 µg/ml), R848 (0.01 µg/ml), GO:R848 (10:0.01) treatment for 24hr. Red staining indicates phalloidin, marker for cell actin showing cell morphology. Nuclei stained as blue with DAPI. Scale bar, 20 µm.

#### Supporting Figure 6

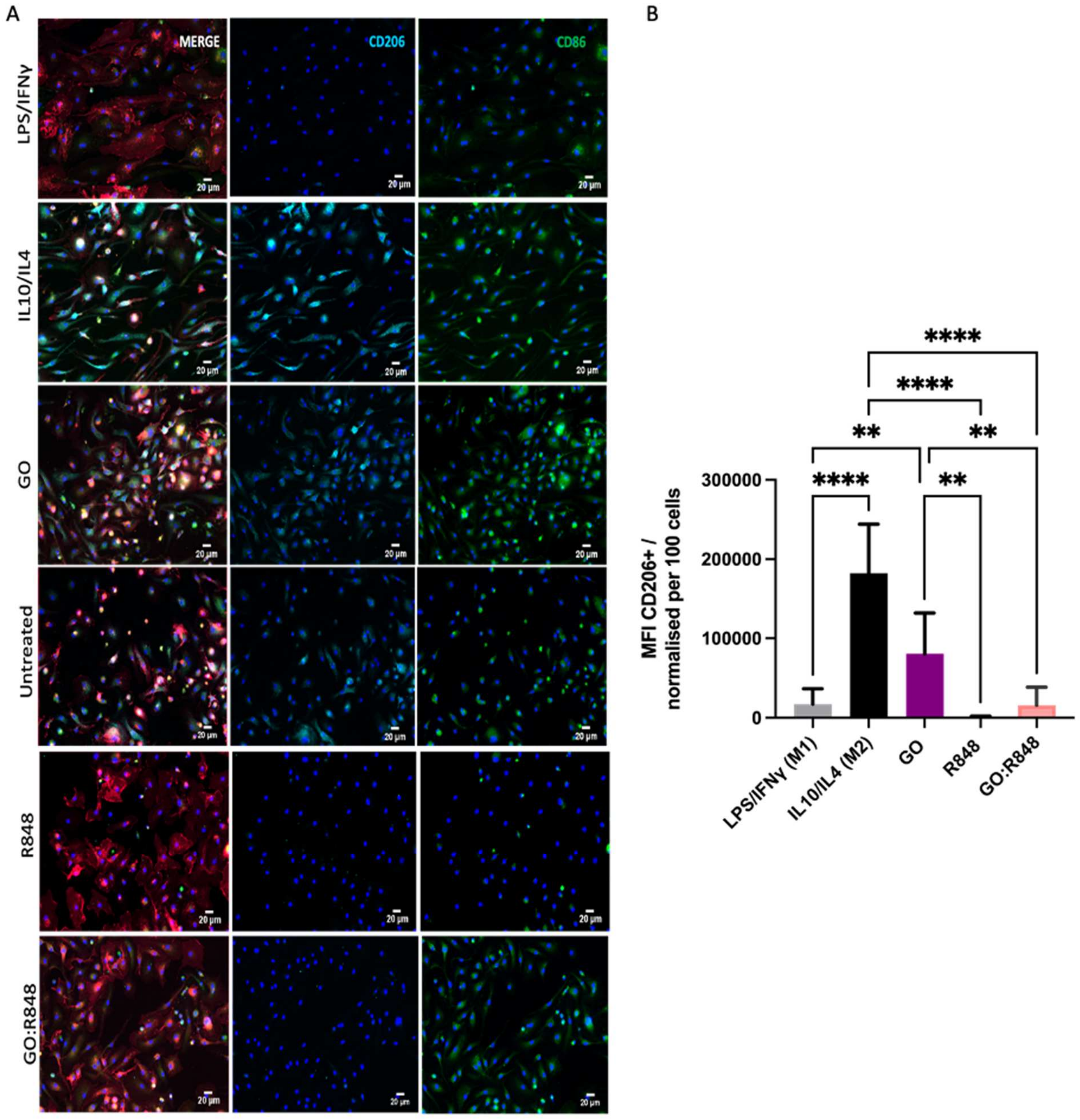


**Figure S6: Reprogramming of M2-like to M1-like BMDMs, 24hr post-treatment.** BMDMs were pre-treated for 24 hr with IL10/IL4 to differentiate them to M2-like, followed by 24 hr of treatment with LPS/IFNγ; M1-like phenotype positive control, IL10/1L4; M2-like phenotype , GO (10 µg/ml), R848 (0.01 µg/ml) and GO:R848 (10:0.01) **A.** Immunofluorescence images 24hr post-treatment with LPS/IFNg, IL4/IL10, GO, untreated, R848 and GO:R848, stained with CD206 (M2 –like marker; cyan), CD80 (M1-like activation marker; green), F480 (Macrophages; red) and DAPI (nuclei; blue). **B.** Mean fluorescence intensity of CD206+ cells normalised by the number of cells per field of view. n=3 biological replicates/condition. Statistical analysis made with one-way ANOVA, Tukey's multi-comparison test. (**p≤0.01, ***p≤0.001, ****p≤0.001).

#### Supporting Figure 7

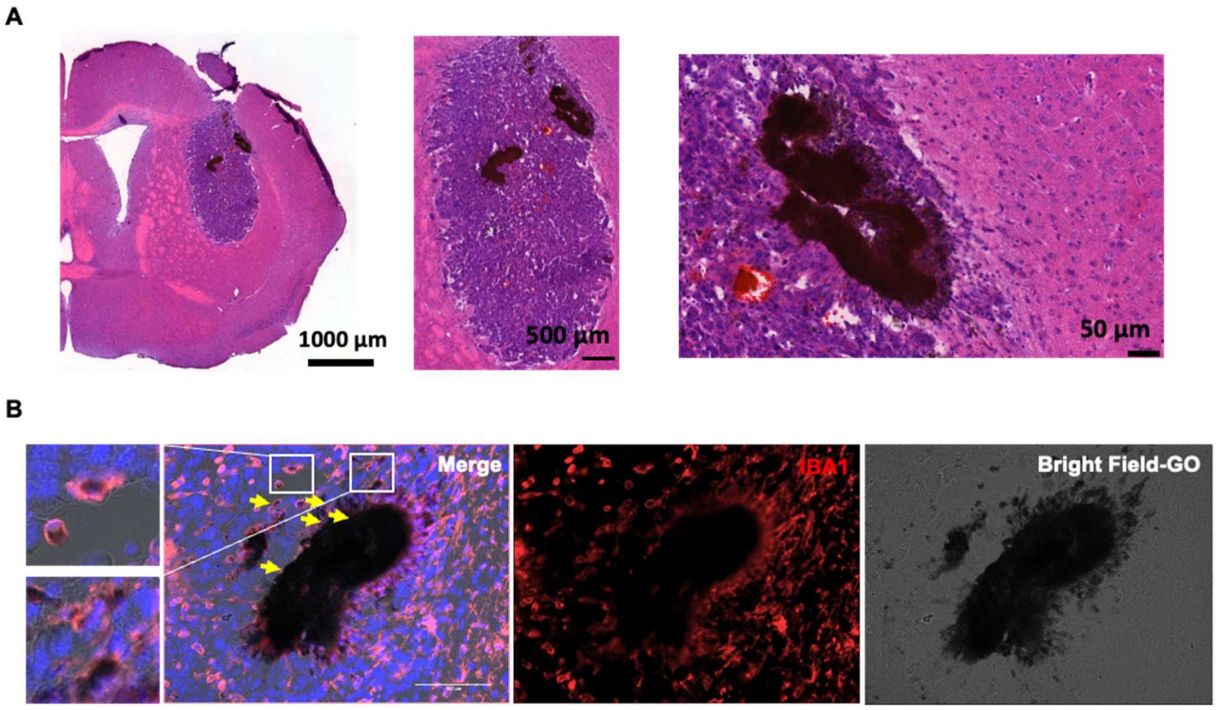


**Figure S7: Graphene oxide conjugated with R848 uptake by TAMMs. A.** Example of GO:R848 spread into GBM tissue 5 days post intratumoral injection. Scale bar, 1000 µm, 500 µm, 50µm. **B**. GO from the GO:R848 complex showed as black (bright field) surrounded by IBA1+ cells (red). Arrows indicates TAMMs full of material driving away from the injection point the particles on day 5 post-treatment. Scale bar 100 µm.

#### Supporting Figure 8

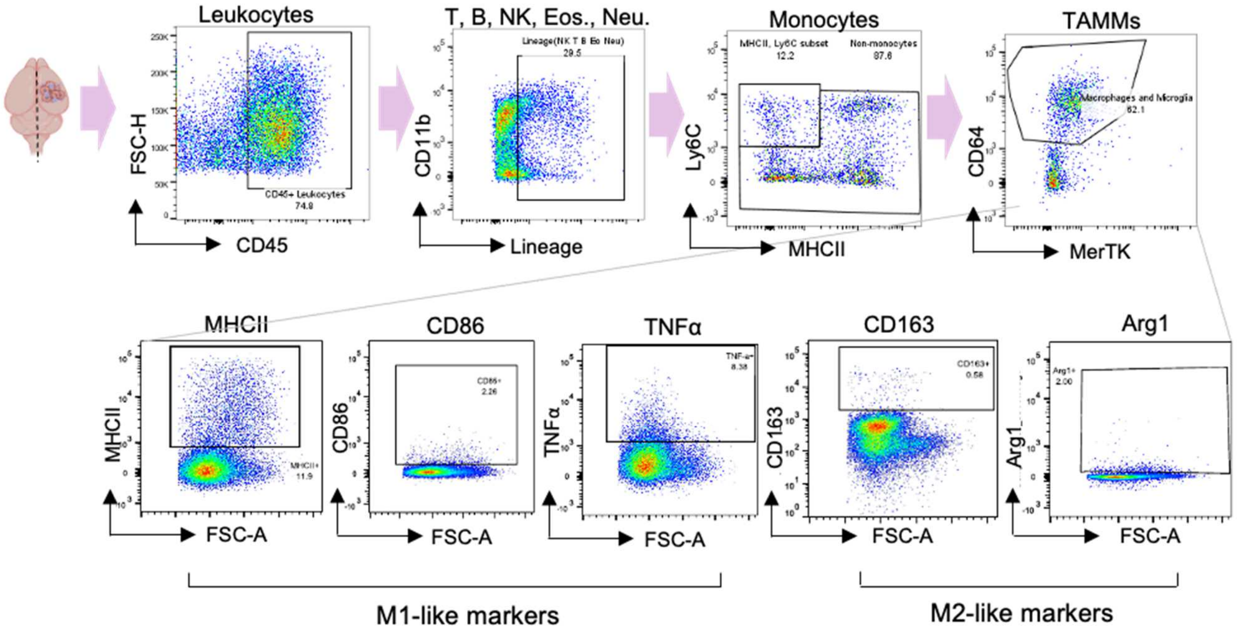


**Figure S8: Example of gating strategy post flow cytometry staining.** Arrows indicate the sequence of gating from live cells gating to microglia and macrophages separation. Lineage cells, eosinophils (Siglec F), neutrophils (Ly6G) and monocytes (Ly6C+/MHCII+) were gated out. M1-like activation markers and pro-inflammatory cytokines (MHCII, CD86, TNF-a), and M2-like pro-tumoral markers (CD163, Arg1) were separated from CD64+/MerTK Monocyte derived-macrophages and microglia cells (TAMMs).

#### Supporting Figure 9

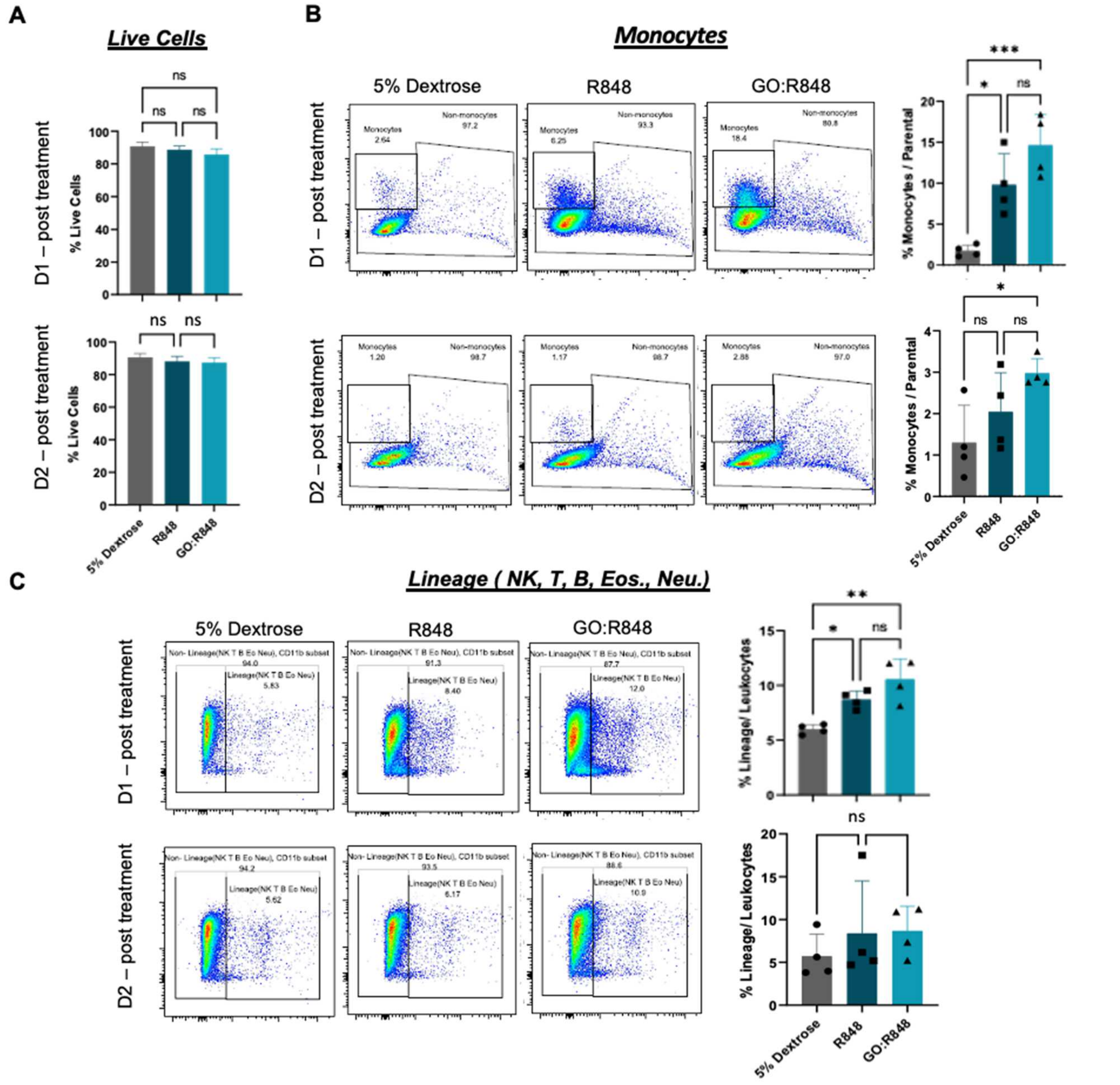


**Figure S9: GO:R848 does not cause cell cytotoxicity and transiently affects lymphocytes and monocytes. A**. Mean of the % of all live cells out of the total population of cells, on day 1 and day 2 post-intratumoral injections of 5% dextrose, R848 and GO:R848 (d5 post-GL261 inoculation). **B.** Percentage of Monocyte/Parental on day 1 and day 2 post-treatment with 5% dextrose, R848 and GO:R848. Representative dot plot flow cytometry graphs showing the gating of monocytes. **C.** Percentage of Lineage cells (Natural killer, B-cells, T-cells, Eosinophils and neutrophils) on day 1 and day 2 post-treatment with 5% dextrose, R848 and GO:R848. Representative dot plot flow cytometry graphs showing the gating of lineage cells per treatments. Statistical analysis made with one-way ANOVA (Tukey's multiple comparison test). Data presented as mean ± SD of n=4 mice/group.

#### Supporting Figure 10

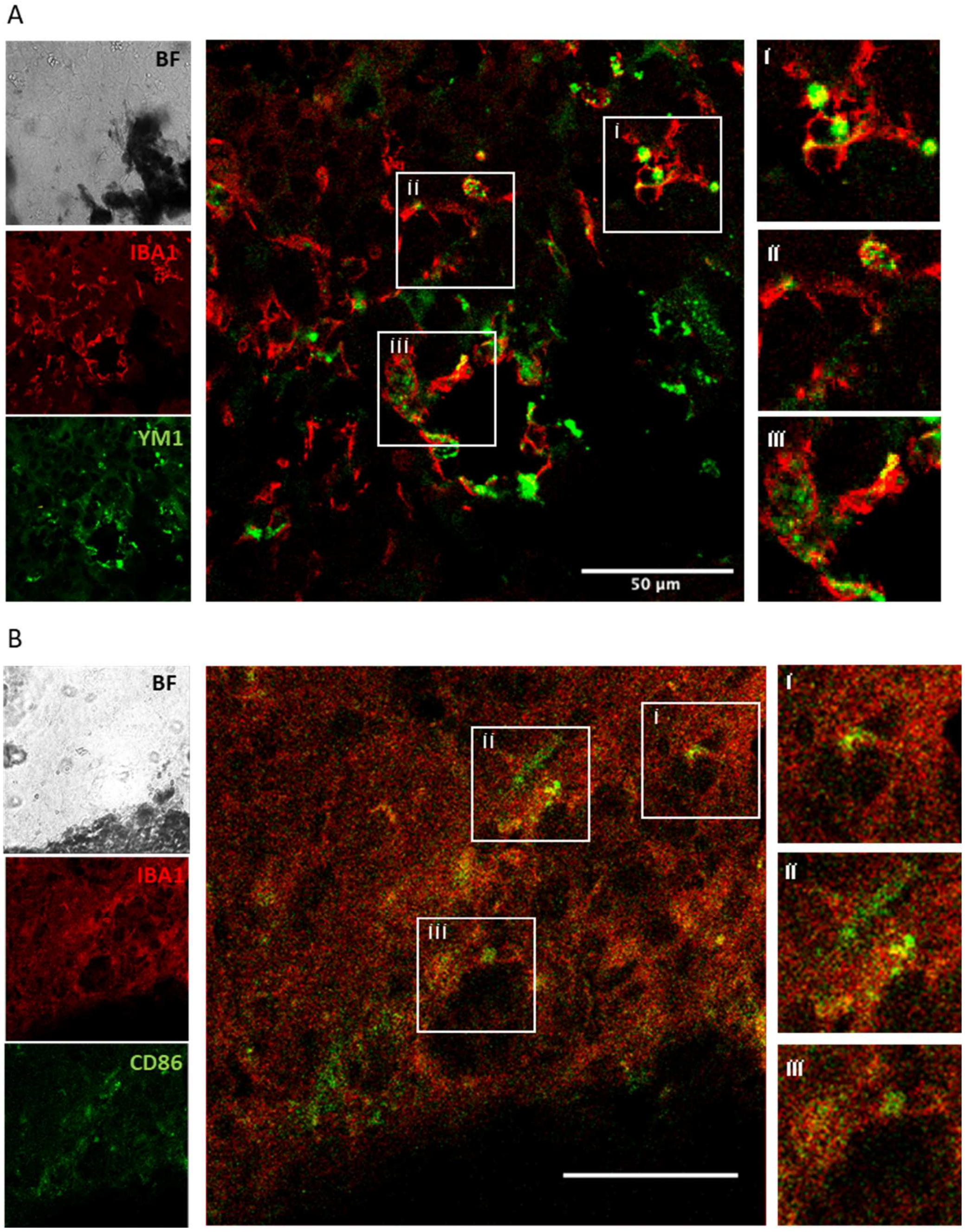


**Figure S10: GO:R848 GO:R848 attracts and is taken up by both M1 and M2 like macrophages .** Representative high magnification confocal images of TAMMs in GBM tissue, 24 hr post GO:R848 injection. A. Images show GO:R848 (bright-field), IBA1+ TAMMs (red), YM1+ M2- like TAMMs (green) and merged image. B. Images show GO:R848 (bright-field), IBA1+ TAMMs (red), CD86+ M1- like TAMMs (green) and merged image. Scale bar, 50 µm. Zoomed in images i, ii, iii.

#### Supporting Figure 11

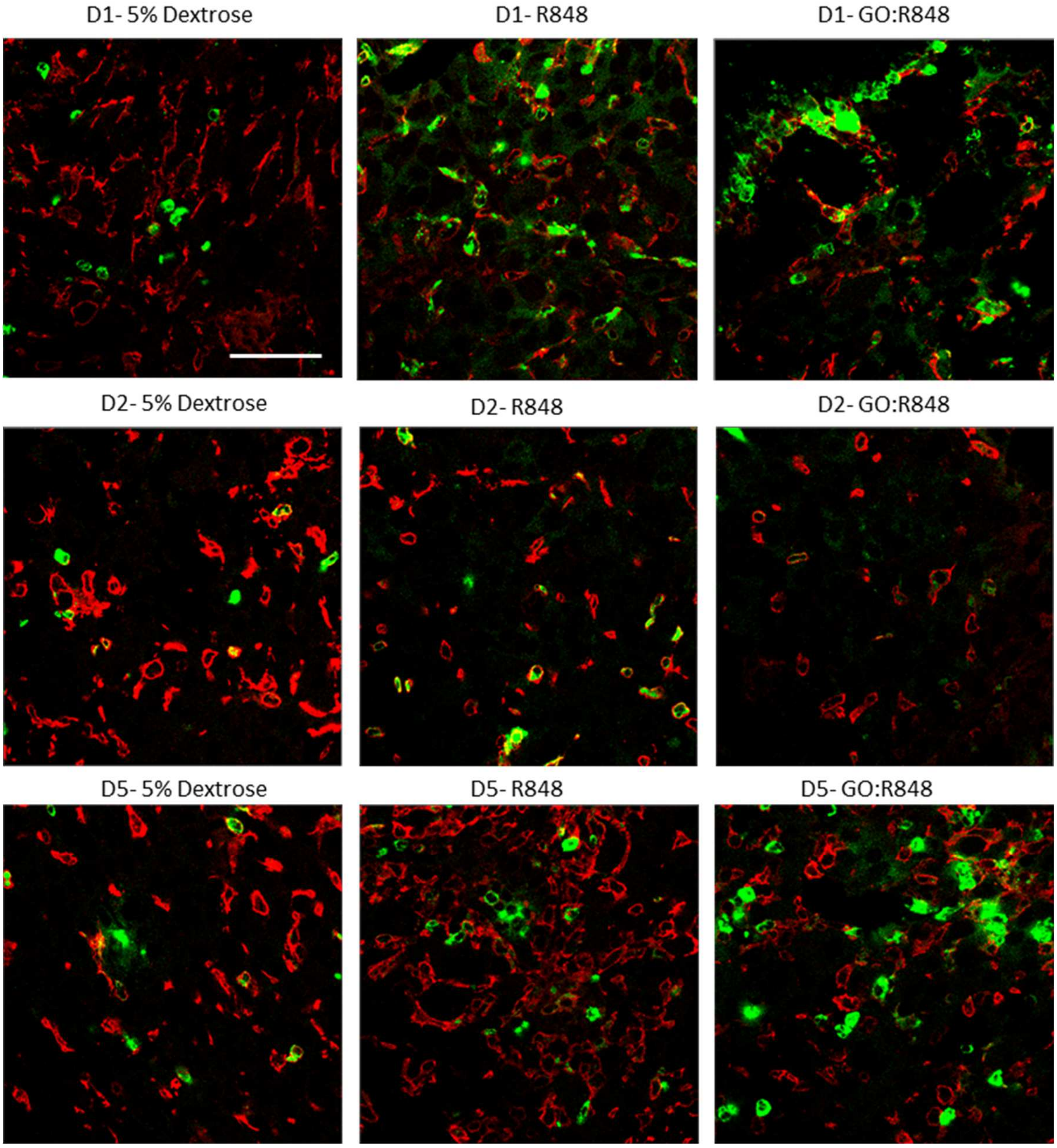


**Figure S11: Elevation of CD86 activation marker and reduction of Ym1 (M2-like) marker.** Representative confocal fluorescent images, of TAMMs within tumour, on day 1, day 2 and day 5 following administration of 5% dextrose, R848 or GO:R848. TAMMs were stained with IBA1+ (red), and CD86 or Ym1 (green). Scale bar, 50 µm.

#### Supporting Figure 12

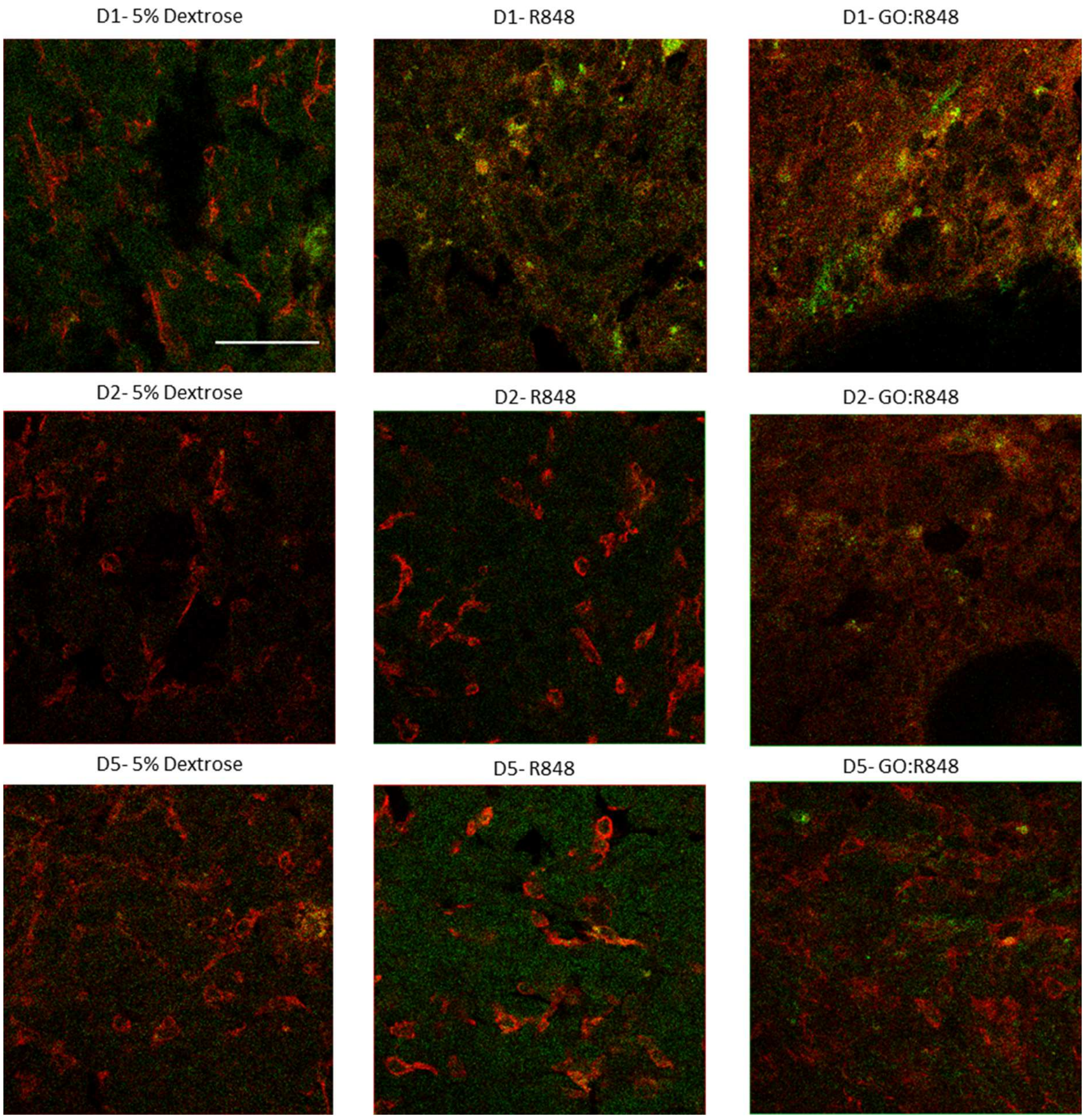


**Figure S12: Elevation of CD86 activation marker and reduction of CD86 activation marker.** Representative confocal fluorescent images, of TAMMs within tumour, on day 1 (D1), day 2 (D2) and day 5 (D5) following administration of 5% dextrose, R848 or GO:R848. TAMMs were stained with IBA1+ (red), and CD86 (green). Scale bar, 50 µm.

### Supporting Figure 13

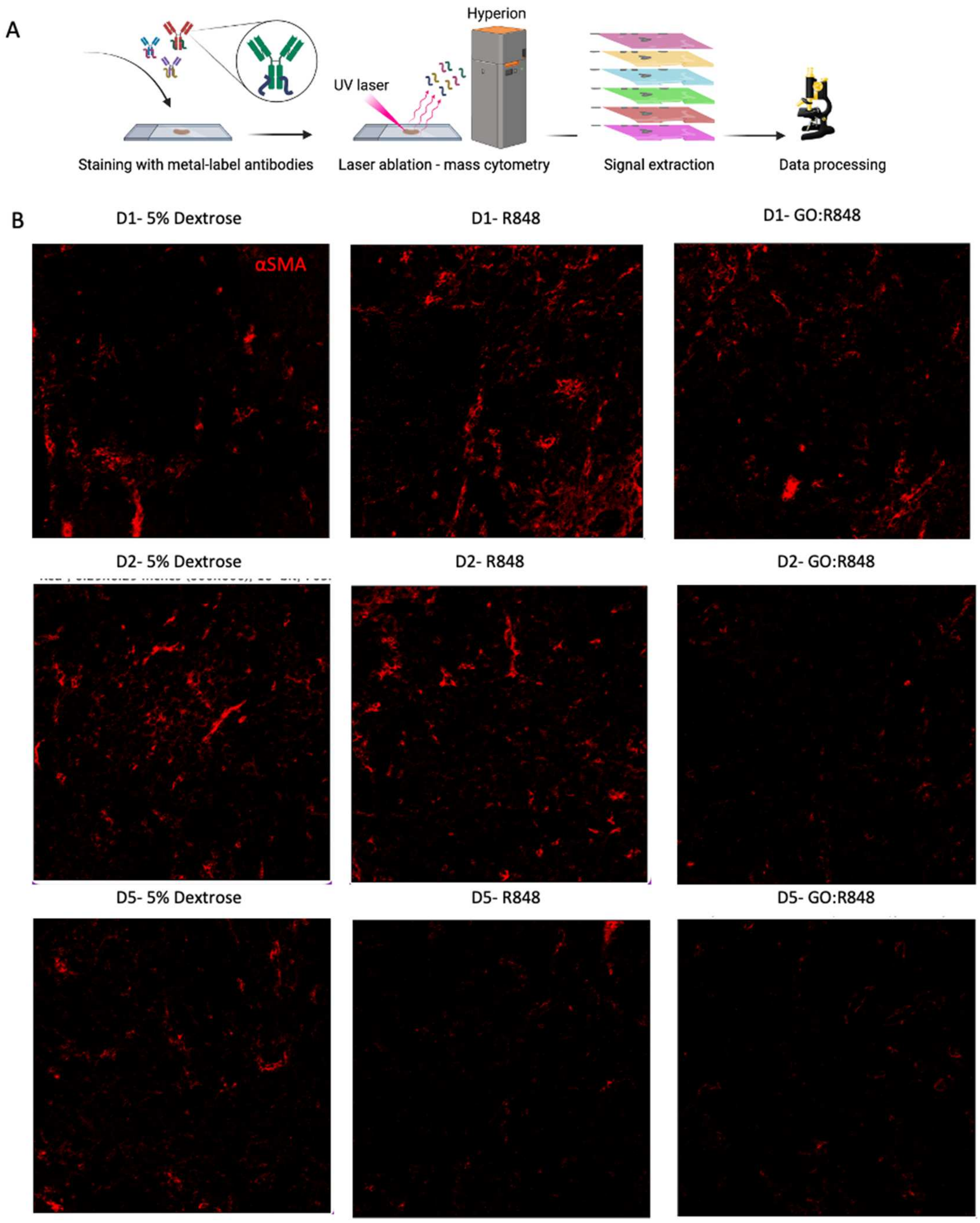


**Figure S13: The effect of GO:R848 on vasculature. A.** Schematic illustration of the experimental set up for image mass cytometry (IMC) using cryo-brain samples, 10µm thickness**. B.** C57Bl/6 mice were implanted with 5x10$^{4}$ (1 µl) GL261-luc cells into the right striatum. Five days following tumour inoculation mice were treated by intratumoral (i.t.) delivery of 5% dextrose, free R848 and GO:R848. Representative images taken by Hyperion (600 µm x 600 µm), presenting vasculature (αSMA; red) on day 1, day 2 and day5 post treatment. Images were normalised based on the negative (unstained) control.

#### Supporting Figure 14

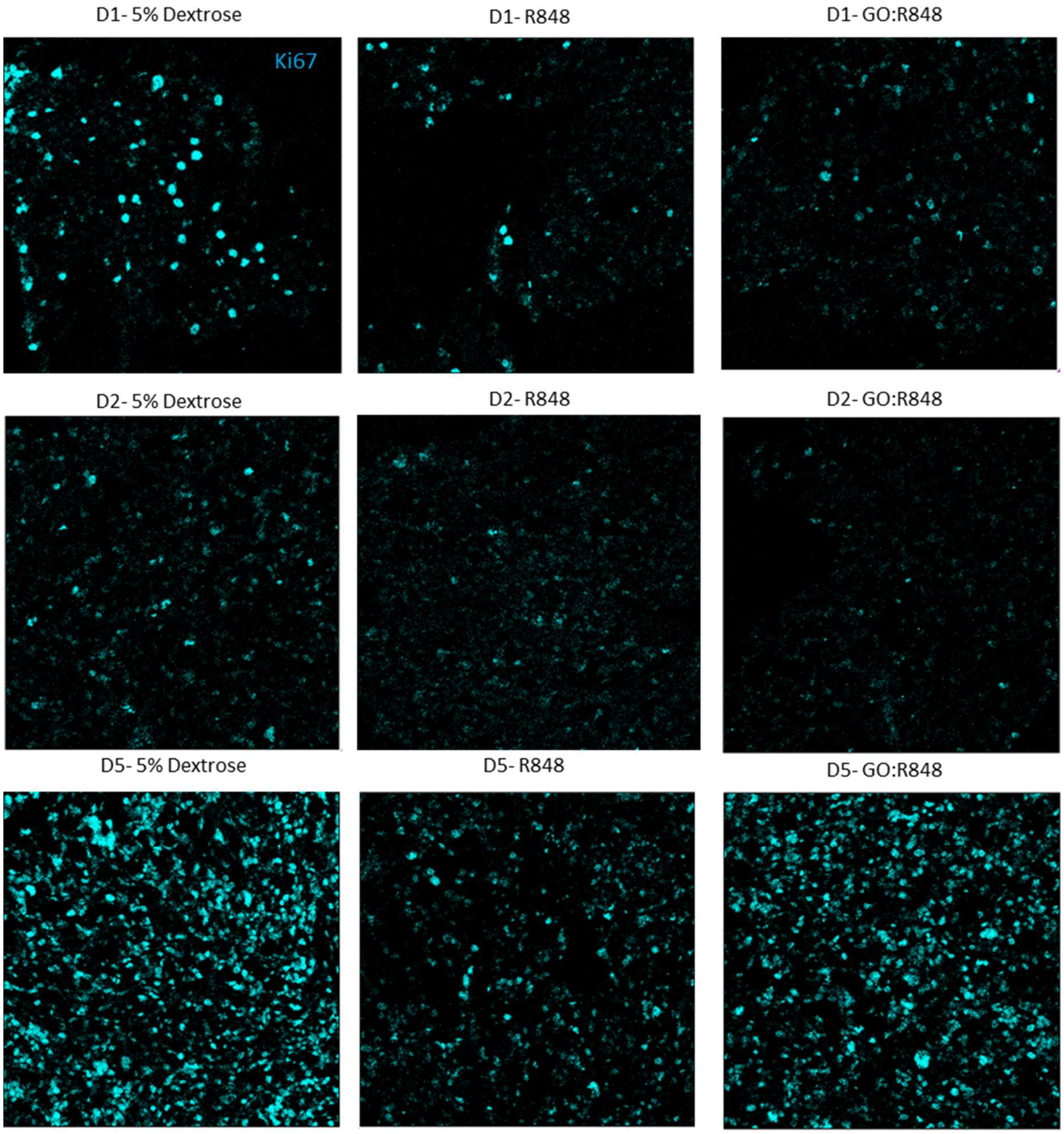


**Figure S14: The effect of GO:R848 on cell proliferation.** C57Bl/6 mice were implanted with 5x10$^4$ (1 µl) GL261-luc cells into the right striatum. Five days following tumour inoculation mice were treated by intratumoral (i.t.) delivery of 5% dextrose, free R848 and GO:R848. Representative images taken by Hyperion (600 µm x 600 µm), showing cell proliferation (Ki67; cyan) on day 1, day 2 and day5 post treatment. Images were normalised based on the negative (unstained) control.

### Supporting Figure 15

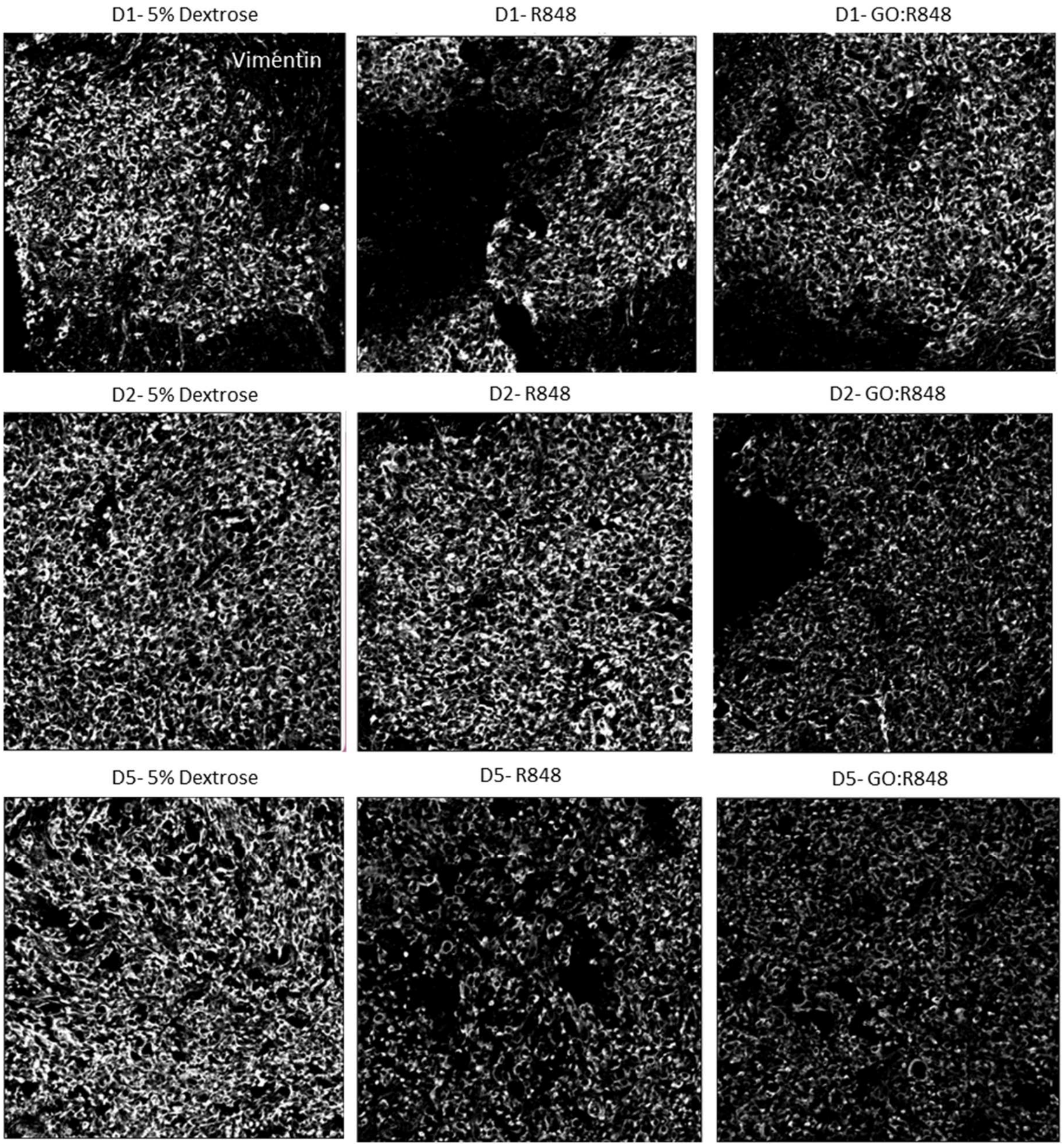


**Figure S15: The effect of GO:R848 on tumour microenvironment.** C57Bl/6 mice were implanted with $5x10^4$ (1 µl) GL261-luc cells into the right striatum. Five days following tumour inoculation mice were treated by intratumoral (i.t.) delivery of 5% dextrose, free R848 and GO:R848. Representative images taken by Hyperion (600 µm x 600 µm), showing Vimentin (grey) on day 1, day 2 and day5 post treatment. Images were normalised based on the negative (unstained) control.

#### Supporting Figure 16

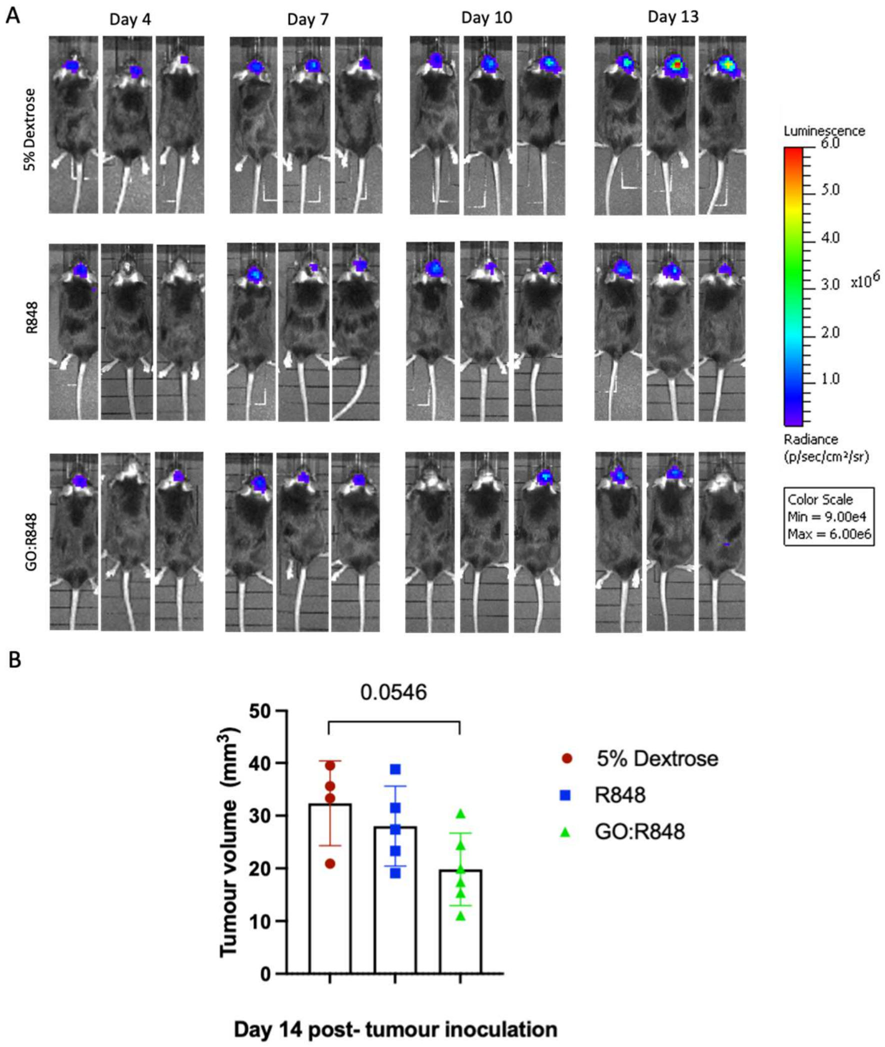


**Figure S16: IVIS imaging pre- and post- GO:R848 treatment and histological analysis showing a delay a of tumour growth. A**. C57Bl/6 mice were implanted with $5x10^4$ (1 μl) GL261-luc cells into the right striatum. BLI was conducted on day 4, as a pre-treatment baseline to normalise mice to different groups. Five- and eight-days following tumour inoculation mice were treated by intratumoral (i.t.) delivery of 5% dextrose (n=7), free R848 (n=6) and GO:R848 (10:4) (n=8). Tumour growth was monitored via BLI on day 4, 7, 10 and 13 post tumour inoculation. In vivo images post-normalisation, on day 4, 7, 10 and 13 of tumour growth. N=3 representative mice per treatment group. **B.** Tumour volume as $mm^3$ calculated on day 14 based on histology for 5% Dextrose, free R848 and GO:R848 treated mice. Data presented as mean ±S.D. One-way ANOVA, Tukey's multi-comparison test.

#### Supporting Figure 17

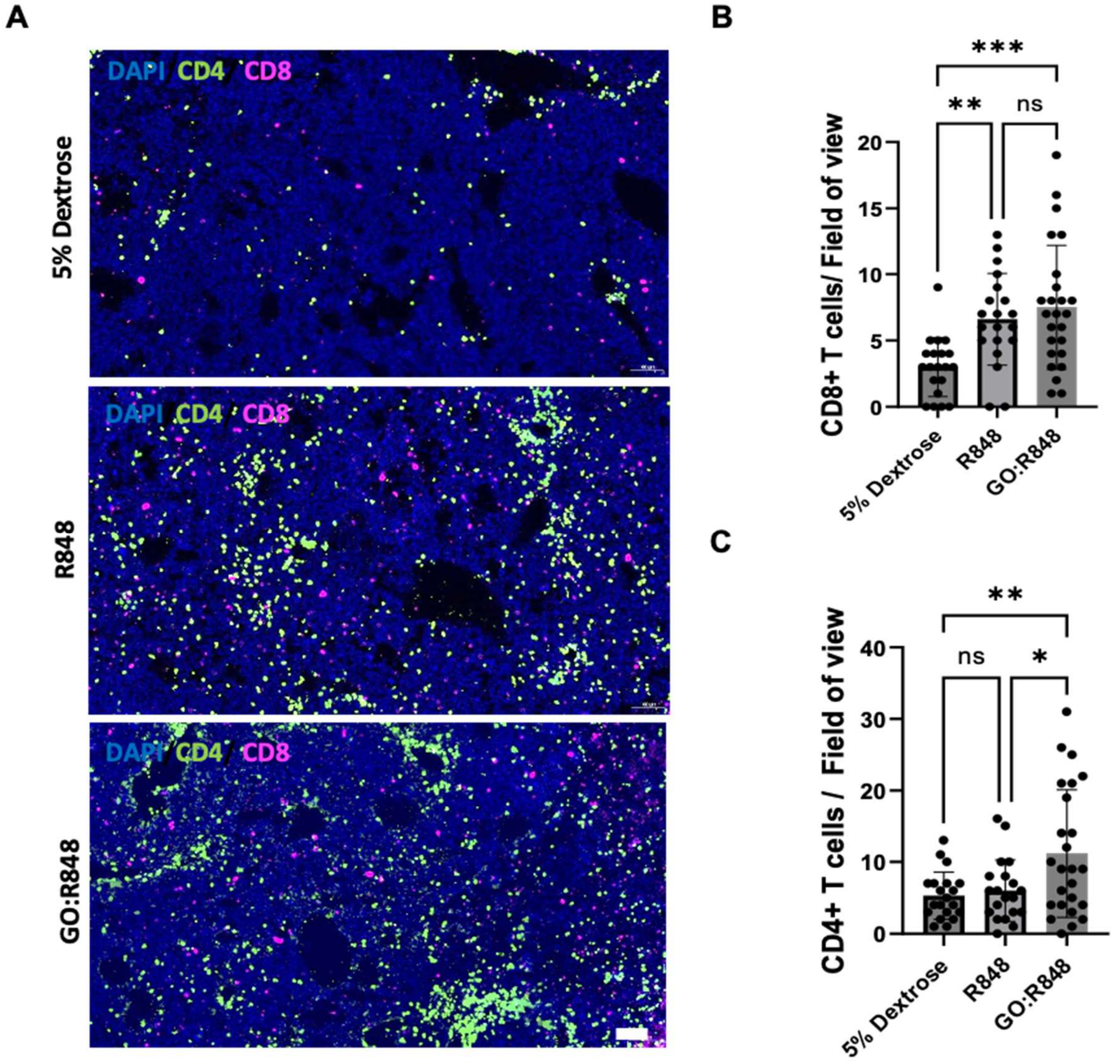


**Figure S17: GO-R848 elevates the recruitment of both CD8+ and CD4+ T cells *in vivo.*** C57Bl/6 mice were implanted with 5x104 (1 µl) GL261-luc cells into the right striatum. Five- and eight-days following tumour inoculation mice were treated by intratumoral (i.t.) delivery of 5% dextrose, R848 (0.72 µg / 3 µl) and GO:R848 (10:4; 1.8 µg : 0.72 µg) (n=4/group) . **A.** Representative slidescanner images showing the recruitment of CD8+ T cells (magenta), CD4+ T cells (green) and nuclei/DAPI (blue), 14 days post tumour inoculation. Scale bar, 100 µm. **B.** Quantification of CD8+ T cells/fields of view and **C.** CD4+ T cells/fields of view on day 14 post-tumour inoculation. Statistical analysis made with two-way ANOVA, Tukey's multi-comparison test. (*p≤0.05, **p≤0.01, ***p≤0.001, ****p≤0.001).
